# A eukaryotic-like phospholipid transfer protein conserved in Asgard archaea

**DOI:** 10.1101/2025.05.16.653879

**Authors:** Nicolas-Frédéric Lipp, Elida Kocharian, Itay Budin

**Affiliations:** Department of Biochemistry & Molecular Biophysics, University of California San Diego, La Jolla, CA 92093, USA

## Abstract

Eukaryotic cells rely on conserved lipid transfer proteins (LTPs) to shuttle insoluble phospholipids between membranes and support organelle proliferation. Because cells lacking organelles do not require extensive lipid transport networks, it is not known if such machinery had to evolve during eukaryogenesis. Here we describe a class of phospholipid transporters in Asgard archaea that share structural and functional features with eukaryotic LTPs of the START domain superfamily. Asgard genomes encode three START protein classes, StarAsg1-3, conserved across most phyla, of which only StarAsg1 has the large hydrophobic cavity and membrane-docking motifs required for lipid transfer. StarAsg1 from Lokiarchaeia binds anionic membranes in vitro and in yeast, extracts phospholipids, and transfers several phospholipid classes between membranes, as eukaryotic START LTPs do. Phylogeny suggests eukaryotic LTPs could share ancestry with StarAsg1, while StarAsg2/3 group with a eukaryotic Hsp90 co-chaperone. Inter-membrane lipid transport may have predated eukaryogenesis and subsequently facilitated organelle proliferation.

---

Eukaryotic organelles rely on proteins that transport lipids from sites of their synthesis, like the endoplasmic reticulum (ER), to organelle membranes. Lipid transfer between membranes is faster than protein trafficking^1,2^ and sustains the high lipid fluxes that organelles require to grow and replicate; it also generates compositional heterogeneities between organelles specialized for their function^3–7^. Intracellular lipid transport occurs via several classes of lipid transfer proteins (LTPs). Some are channel-shaped proteins that form bridges across contact sites, where organelles sit within tens of nanometers from each other^8,9^. However, most are small, soluble proteins that transiently interact with membranes and transport individual lipid molecules between membranes^10,11^. These shuttle-like LTPs rely on soluble domains with amphipathic membrane-binding sites, regulated by dynamic bound-to-soluble switching mechanisms that support second-to-minute-scale lipid transport rates^12,13^. Despite their central roles in the architecture of eukaryotic cells, models for the origin of eukaryotic LTPs remain limited.

Eukaryotes arose from archaea between 1.5 and 2.5 billion years ago^14–17^, with mitochondria descended from an engulfed proto-symbiont sister to alphaproteobacteria^18–20^. In bacteria, phospholipid transfer can occur across the periplasmic space through LolA/B and MlaC homologs^21^,but no direct eukaryotic counterparts to these systems have been identified. In contrast, little is known about any LTPs in archaea. Promethearchaeati^22^ (hereafter called Asgards) are an archaeal superphylum that forms a monophyletic group with Eukarya^23^ and recent analyses suggest eukaryotes belong to Asgards *sensu stricto*^24–26^. Asgards contain multiple Eukaryotic Signature Proteins (ESPs), including homologs of key components of eukaryotic membrane machinery^27–31^. Cultured Asgard, like the Lokiarchaeon *Promethearchaeum syntrophicum* MK-D1^22,32^, show complex membrane topologies including tubular protrusions. Emerging molecular clock data support a late-mitochondrial model for eukaryogenesis, in which the Asgard ancestor was already quite complex^33^. Recent imaging of exponential phase Heimdallarchaeia revealed large, intracellular vesicles^34^ that resemble eukaryotic compartments like autophagosomes that rely on LTPs for their formation and growth^35^. The presence of internal compartments in Asgard cells implies a need for inter-membrane lipid transport that could have been co-opted for organelle proliferation during eukaryogenesis. We therefore asked whether extant Asgards encode LTPs related to those of eukaryotes.

Here we describe three proteins in the Steroidogenic Acute Regulatory protein-related lipid Transfer (START)-domain family in Lokiarchaeia that are widely conserved across Asgard lineages. In eukaryotes, START domains represent a widely utilized protein fold for LTPs, but can also function as regulatory proteins^36–38^, transcription factors^39^, and co-chaperones^40^. We report that one Lokiarchaeia START protein is specialized for interaction with membranes and transports phospholipids *in vitro*, similarly to other START domain LTPs. Several lines of evidence suggest a connection between this protein family and eukaryotic START LTPs, while another Asgard START domain protein is ancestral to a conserved eukaryotic co-chaperone.

## Results

### START domain proteins in Lokiarchaeia

We first examined the distribution of 14 annotated classes of shuttle-like LTPs in bacteria, archaea, viruses and eukaryotes (**Extended Data Fig. 1**). Nine of them are exclusive to eukaryotes with very rare occurrences in other groups, suggesting that the diversity of LTPs expanded during eukaryotic evolution. In contrast, proteins containing START and Sterol Carrier Protein 2 (SCP-2) domains are distributed among eukaryotes and archaea, but also found in bacteria. From this observation, we identified one SCP-2 homolog and three START-like domains in the isolated Lokiarchaeon *P. syntrophicum* MK-D1, which we named StarLok1, StarLok2 and StarLok3. Although we did not observe any clustering of nearby genes related to lipid metabolism, we focused attention on these candidates because the structural features and ability to bind hydrophobic ligands of START domains are well defined, in contrast to other families like SCP-2^41,42^.

We further characterized the predicted structural features in the three MK-D1 START domains. We observed a charge distribution of StarLok1 that is strongly polarized, with a large cationic patch surrounding the mouth of the cavity (**Fig. 1a**). This configuration suggested that StarLok1 may interact with membranes, as peripheral membrane proteins are often electrostatically polarized to interact with phospholipids^43–46^. A network of hydrophobic residues (F20, F23, L26, W29, F77, V90, W104, F110, F138), two opposite tyrosines (Y54, Y81), and two distant anionic residues (D21, E66) characterize the cavity of StarLok1, supporting the binding of hydrophobic ligands. Notably, the cavity lacks key polar residues generally required for enzymatic activity, such as ketone cyclization^47^. The Positioning of Protein in Membrane (PPM) algorithm suggested that StarLok1 could dock to membranes through the insertion of five hydrophobic residues, reminiscent of the binding mode of the sterol transporter STARD4^48^ (**Fig. 1b**). In contrast, StarLok2 and StarLok3 have lower electrostatic polarization, minimal cavities, and no clear predicted mode of membrane interaction by PPM. To test the accuracy of structural predictions, we expressed and purified the three proteins from *Escherichia coli* (**Supplementary Fig. 1a-e**). Folding was confirmed by Circular Dichroism (CD) spectroscopy (**Fig. 1c**), which showed a secondary structure profile similar to those of previously profiled START domain homologs^49,50^. Among the three, StarLok1 showed reduced ellipticity, StarLok2 featured an increased helical signal and reduced β-strand proportion, and StarLok3 showed a reduced number of turns. These features and the overall spectra matched those derived from the AlphaFold structures (**Supplementary Fig. 1f-h**). Overall, structure-sequence analyses identified the Bet v 1 fold as the closest experimentally determined structural template for StarLok1, reflecting the shared helix-grip scaffold, while StarLok3 resembled human AHA1 START domain (AHSA1) **(Fig. 1d, e)**.

**Fig. 1.**
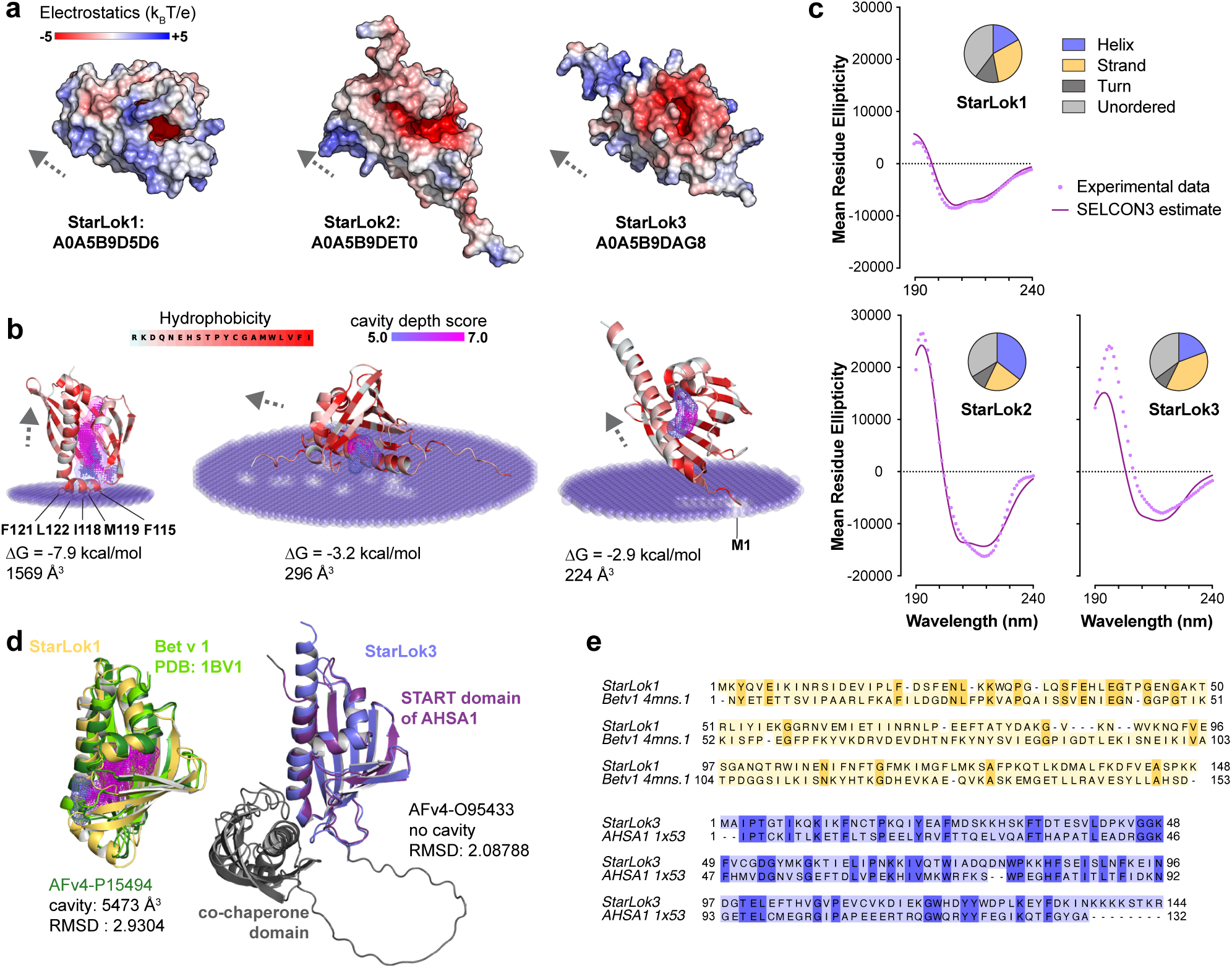
START domain proteins from Lokiarchaea have divergent structural features. **a.** Structural prediction of StarLok1, StarLok2 and StarLok3, represented as electrostatic surface potential. Arrows indicate the orientation of the main α-helix of each START domain, from N- to C-terminus. **b.** PPM prediction of StarLok1, 2 and 3 docked to a model archaeal lipid membrane. The structure is colored according to the hydrophobicity of each residue from pale cyan (low hydrophobicity) to red (high hydrophobicity). Computed cavities are visualized as a 3D point cloud, color-coded by LigSite depth scores, from periwinkle blue (lower scores) to magenta (higher scores). Protein-membrane interaction energies and computed cavity sizes are provided for each protein. **c.** CD analysis of StarLok1, 2 and 3. Experimental mean residue ellipticity (MRE in deg.cm^2^.dmol^-1^) was recorded from 190 to 240 nm (dotted lines) and secondary structures were assigned by SELCON3 decomposition (solid lines). Pie charts summarize the estimated proportion of α-helix, β-strand, turn and unordered regions for each protein. **d.** *Left:* Structural similarities between the AF structure of StarLok1 (yellow) and that of Bet v 1 (dark green). The crystal structure of Bet v 1 (pdb: 1BV1, light green) was also aligned. The cavity of Bet v 1 is displayed as in panel A using LigSite depth score. *Right panel:* Structural similarities between the AF structure of StarLok3 (blue) and that of the START domain of AHSA1 (Purple). The co-chaperone binding domain of AHSA1 is indicated in black. RMSD of AFv4-P15494 and AFv4-O95433 aligned to StarLok1 and StarLok3, respectively, are indicated. The cavity of AFv4-P15494 is also reported. **e.** SWISS-MODEL hit template alignment of StarLok1 and Bet v 1, and StarLok3 and AHSA1 sequences.

### Distribution of START domains across Asgard archaea

To characterize the phylogeny of START domains within Asgards, we analyzed the relationships of 55 homologous sequences of this class across phyla. The resulting tree indicates that StarLok1 is part of a monophyletic clade comprising 21 sequences, with moderate SH-aLRT support and limited UFBoot support (SH-aLRT/UFBoot = 91.4/61) (**Fig. 2a**), a placement we treated as a hypothesis tested below. Proteins of this clade (‘StarAsg1’) feature predicted binding cavities large enough (MK-D1 1569 Å^3^; Heimdall AB_125, 1579 Å^3^) to accommodate a wide range of lipids, including phospholipids (∼1,000 Å^3^) (**Fig. 2b**), consistent with a lipid-binding function. In contrast, the clades containing StarLok2 (StarAsg2) and StarLok3 (StarAsg3) have smaller binding cavities, with StarAsg3 homologs possessing residual pockets under 300 Å^3^. StarLok2 and StarLok3 proteins, but not StarAsg1, were annotated as homologs of the activators of the heat shock protein 90 (Hsp90) ATPase (AHA1) by the database-derived Protein Natural Language Model (ProtNLM/ EMBL-EBI)^51^.

**Fig. 2.**
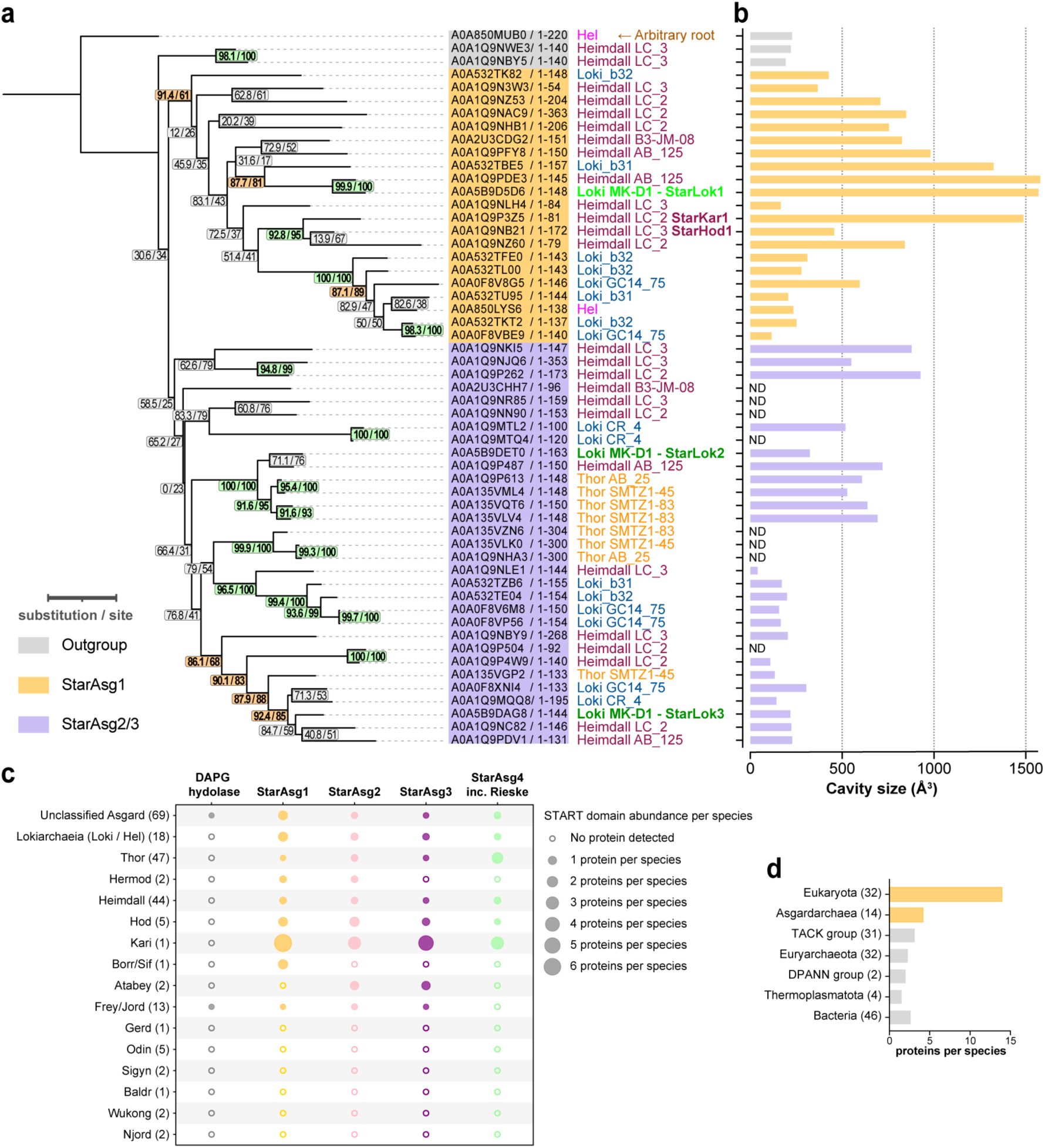
Distinct clades of START domain proteins across Asgards. **a.** ML phylogenetic tree (PhyML Q.Pfam + R4 + F) of the START proteins identified in Asgards. Branch support values are indicated as SH-aLRT (%) and ultrafast bootstrap (%). Supports ≥ 90/90 are highlighted in green, while supports ≥ 85/60 are indicated in orange. The tree is arbitrarily rooted with a single sequence identified in the Helarchaeales (strain CR_Bin_291). Species names are color-coded according to their taxonomic groups: Helarchaeales (magenta), Heimdallarchaeia (wine), Thorarchaeia (orange), Lokiarchaeia (blue), except for MK-D1 (green). Sequence accessions and lengths corresponding to the monophyletic clade of StarAsg1 are highlighted in yellow, while the monophyletic clade of StarAsg2 and StarAsg3 is highlighted in purple. **b.** Comparative analysis of cavity sizes in StarAsg1 and its homologs in Asgards. The UniProt accession numbers are listed along the y-axis, with the corresponding cavity sizes represented by horizontal bars. Yellow bars indicate proteins belonging to the StarAsg1 clade, and purple bars indicate proteins from the StarAsg2 and StarAsg3 clades. ND indicates the absence of a detectable cavity. **c.** Abundance of the START domain classes in different Asgard phyla. The dot size is proportional to the number of identified START domains per species; an empty circle indicates the absence of START domains in that phylum. The average number of determined species in the dataset is given next to each phylum’s name in parentheses. **d.** Abundance of the START domains identified in a representative set of archaeal, bacterial and eukaryotic species referenced in the UniProt database. The number of species in each clade is indicated in parentheses. The set corresponds to that used for the analyses in Fig. 5.

In a holistic analysis of Asgard proteomes using InterProScan profile signatures **(Methods)**, we identified 411 START domain sequences distributed across 203 extant species. We did not observe any START homolog in several phyla, including Odin and Wukong. However, the three START domain clades (StarAsg1-3) are well represented in most phyla with Kari, Hod and Heimdall encoding all three. We also identified three occurrences of DAPG hydrolase (PFAM 18089) as well as twenty-nine Rieske [2Fe-2S] type proteins diverging from a group of START domains, which we named StarAsg4 (**Fig. 2c**). This analysis indicated a distinction between StarAsg1 (SH-aLRT/UFBoot = 83.3/83), StarAsg3 (73/81) and the group comprising StarAsg4 and StarAsg2 (86.1/71) (**Extended Data Fig. 2**), but these hypotheses required additional analyses given their limited branch support. We thus sought to further determine the divergent structural and functional features of these proteins through experiments.

### StarLok1 interacts with anionic membranes

Our analyses suggested StarLok1 could be uniquely suited for membrane binding. Consistent with this model, StarLok2 and StarLok3 showed high solubility, but StarLok1 aggregated when concentrated in the absence of detergents. Detergent-free StarLok1 could be refolded and stabilized at acidic pH, which promotes increased cationic electrostatic repulsion along the amphipathic protein face, and was additionally stabilized by acetate (**Supplementary Fig. 2**). StarHod1, an ortholog of StarLok1 present in Hodarchaeales, showed similar predicted structural features and stabilization by acetate (**Supplementary Fig. 3**). Lokiarchaeia feature an acetogenic metabolism^52^ that releases acetate as a byproduct of ATP production^53^, which could thus influence StarLok1 solubility and interactions.

We assayed the interaction of each Lokiarchaeia START protein with membranes. Addition of stabilized and folded StarLok1 (preparation described in **Supplementary Fig. 4**) triggered clustering of liposomes containing 5% or higher concentrations of phosphatidylserine (PS), an anionic phospholipid, but not in neutral di-oleoyl-phosphatidylcholine (DOPC) liposomes (**Fig. 3a**). In contrast, StarLok2 and StarLok3 showed minimal interactions with PS-containing liposomes. We also detected gravity separation of StarLok1 in solutions with PS-containing liposomes, but not DOPC liposomes (**Supplementary Fig. 4g**). StarLok1 displayed other features consistent with its hydrophobic nature, including self-quenching of hydrophobic dyes on SDS-PAGE (**Supplementary Fig. 4c**), and accumulation at air-water interfaces (**Supplementary Fig. 4g**). When StarLok1 was expressed as a fusion protein to GFP in *Saccharomyces cerevisiae*, it showed strong plasma membrane (PM) localization, with additional enrichment at the bud neck (**Fig. 3b**). StarHod1 showed a similar distribution, co-localizing with an established PS-binding sensor, the C2 domain of lactadherin (C2_LACT_)^54^ (**Fig. 3c**). In contrast, StarLok2 and StarLok3 remained soluble in the yeast cytosol (**Fig. 3d**).

**Fig. 3.**
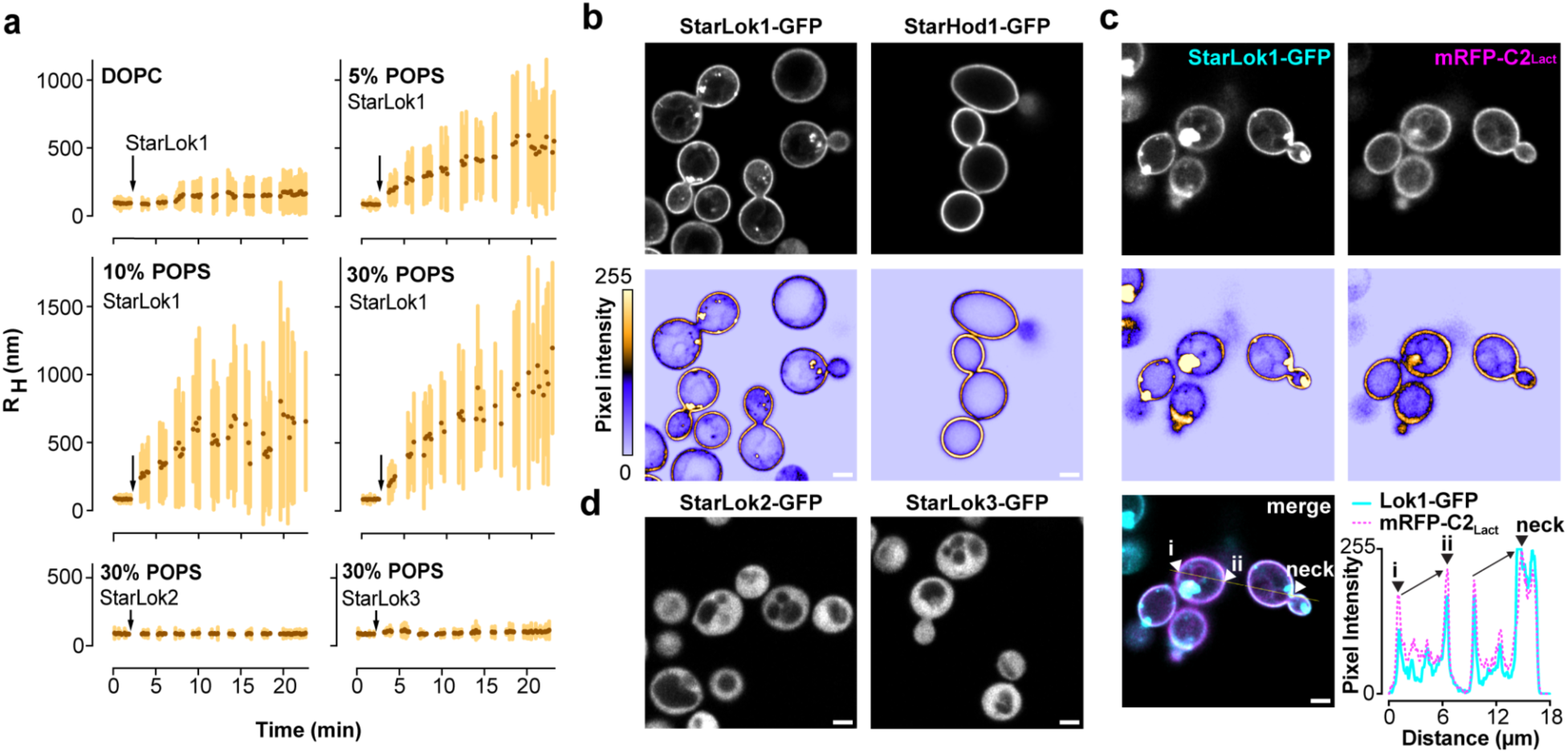
StarAsg1 proteins are specialized for interaction with anionic membranes. **a.** DLS of StarAsg proteins with liposomes of different lipid compositions. Light scattering was recorded for 2 min after the addition of 50 *µ*M liposomes composed of pure DOPC, or DOPC doped with 5 mol% POPS, 10 mol% POPS, or 30 mol% POPS. At t = 2 min, 1 *µ*M StarLok1 was added to the mixture. Increasing POPS causes protein-mediated aggregation of the liposomes. Addition of StarLok2 or StarLok3 to DOPC liposomes with 30 mol% POPS does not induce aggregation. **b.** StarLok1-GFP and StarHod1-GFP expressed in budding yeast bind the PM. The lower panels are intensity heat maps of the corresponding confocal image to highlight the polarized distribution of StarLok1 and StarHod1 in membranes surrounding the yeast bud neck and budding daughter cells. **c.** StarLok1 localization and polarization correlate with PS abundance in the PM. PS is detected with the protein sensor mRFP-C2_LACT_. Composite image shows StarLok1-GFP (cyan) and mRFP-C2_LACT_ (magenta), both of whose signal increases along the PM towards the bud neck, indicated in the plot profile. Scale bars, 2 *µ*m.

### StarLok1 binds phospholipids and transfers them between membranes

We sought to identify bound lipid ligands^55^ of Asgard LTPs expressed in cells by isolating GFP-tagged StarLok1, 2, and 3 and StarHod1 from yeast cells and analyzing their bound lipids by LC/MS^2^ (**Fig. 4a**). While StarLok2 and StarLok3 showed low lipid binding, StarLok1 and StarHod1 extracts contained bound phospholipid (∼110 pmol) roughly equimolar to the protein-binding capacity of the anti-GFP beads (∼150 pmol). StarLok1-bound ligands were notably enriched in PS, while StarHod1 ligands were enriched in phosphatidylcholine (PC) (**Fig. 4b**). Both also bound phosphatidylethanolamine (PE). StarLok2 and StarLok3 extracts contained fewer lipids, which were predominantly phosphatidylinositol (PI) and PC, likely reflecting background binding of the major yeast whole-cell phospholipids^56^.

**Fig. 4.**
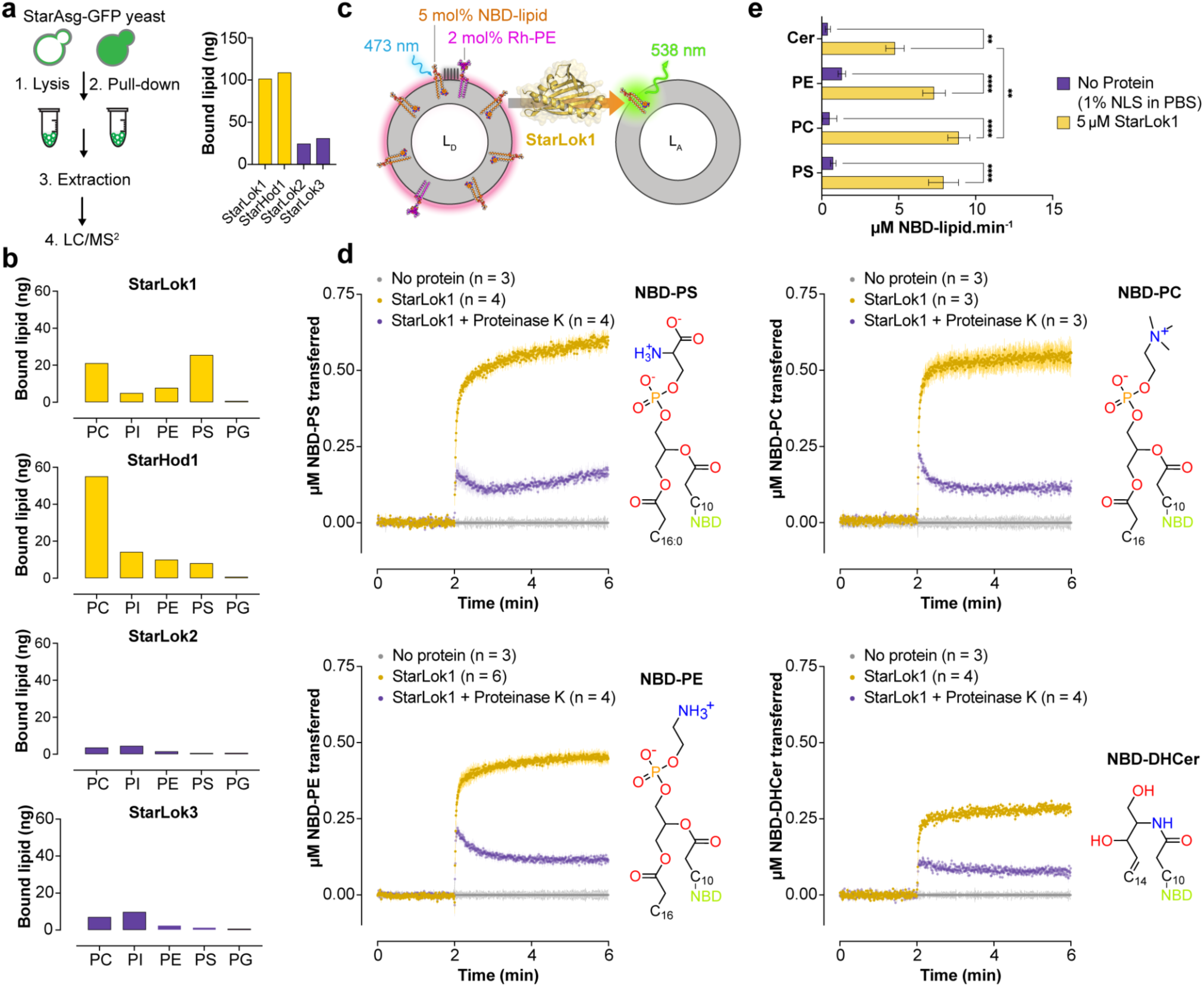
Characterization of lipids bound to StarAsg proteins and lipid transfer activity. **a.** Schematic representation of the experimental workflow used to identify lipids bound to GFP-tagged StarAsg proteins expressed in yeast and plot showing total lipid abundance quantified by LC/MS^2^. **b.** Abundance of phospholipid classes associated with StarLok1, StarHod1, and StarLok2/3. StarAsg1 proteins show increased levels of PC, PE, and PS binding. **c**. Schematic of lipid transfer assay. Donor liposomes (L_D_, 200 µM total lipid) and acceptor liposomes (L_A_, 200 µM total lipid) were incubated with 5µM of StarLok1 in PBS buffer (pH 7.4). **d.** NBD-lipid transfer kinetics of StarLok1 (yellow) incubated with L_D_ and L_A_ liposomes compared to a blank control containing an equivalent amount of N-lauroylsarcosine (gray), or an equivalent molar amount of proteinase K–inactivated StarLok1 (purple). The fluorescence signal was normalized using calibration control to yield the absolute amount of NBD-lipid transferred (µM) and is represented as the mean of n independent replicates ± SD. Each plot represents a different NBD-tagged ligand, whose structure is shown to the right. **e.** Calculated lipid transfer rate derived from the initial slopes of the transfer curves (mean ± SEM). Statistical significance was determined using two-way ANOVA followed by Tukey’s multiple comparisons test. \*\**P*< 0.01; \*\*\*\**P*< 0.0001.

Based on its capacity to bind phospholipids, we asked if StarLok1 could act as a *bona fide* LTP. The movement of NBD-labeled phospholipid analogues from donor to acceptor liposomes can be monitored using a Förster resonance energy transfer (FRET)-based assay in which Rhodamine-PE acts as a quencher bound to donor liposomes (**Fig. 4c**), as previously shown for STARD2, STARD10, PITPα/β and CERT^57–59^. Upon addition of detergent-solubilized StarLok1 to donor liposomes containing NBD-PS, NBD-PC, or NBD-PE, a robust increase in NBD fluorescence was observed, indicating lipid transfer (**Fig. 4d**). Phospholipid transport was not observed beyond baseline when equivalent detergent was added, or when the protein was digested with Proteinase K (**Supplementary Fig. 5a**). Using equilibrated liposomes, we quantified initial transport rates ranging between 1-2 phospholipid·min^-1^ per StarLok1 (**Fig. 4e**). Transport of NBD-labeled ceramide, a lipid that lacks a phosphate headgroup, was slower than that of the phospholipids, while transport of Rhodamine-PE itself, which contains a very large headgroup, was not detected in control liposomes pre-equilibrated for NBD-lipid (**Supplementary Fig. 5b, c**).

### Sequence and structural phylogeny of START domains across the tree of life

Motivated by our biochemical data, we sought to understand the phylogenetic relationships across START domains and test the likelihood that eukaryotic LTPs originated from Asgards. We applied two complementary approaches, sequence based maximum-likelihood (ML) and structural phylogeny, to a manually curated set of 161 representative taxa containing eukaryotes, bacteria, and archaea across the tree of life including non-Asgard archaeal lineages Euryarchaeota, TACK group, DPANN group, and Thermoplasmatota. The full dataset comprised 774 START domain-containing sequences. For the ML approach, we screened several sequence alignment algorithms to optimize the overall conservation score of different sequence subsets and generated several ML trees (Fig. 5a, **Supplementary Figs. 6-12**) to address specific questions. For both approaches, and because START domains often exist in tandem repeats or as parts of multi-domain proteins, we developed a tool (ProtDomRetriever) to isolate individual domains, yielding 842 domains across the 774 sequences (**Fig. 5b**). For structural phylogeny, we employed the FoldTree^60^ algorithm, which generates trees based on structural divergences measured by Foldseek^61^, applied to the 805 domains recovered from the 748 sequences for which AlphaFold structures were available **(Fig. 5c).**

**Fig. 5.**
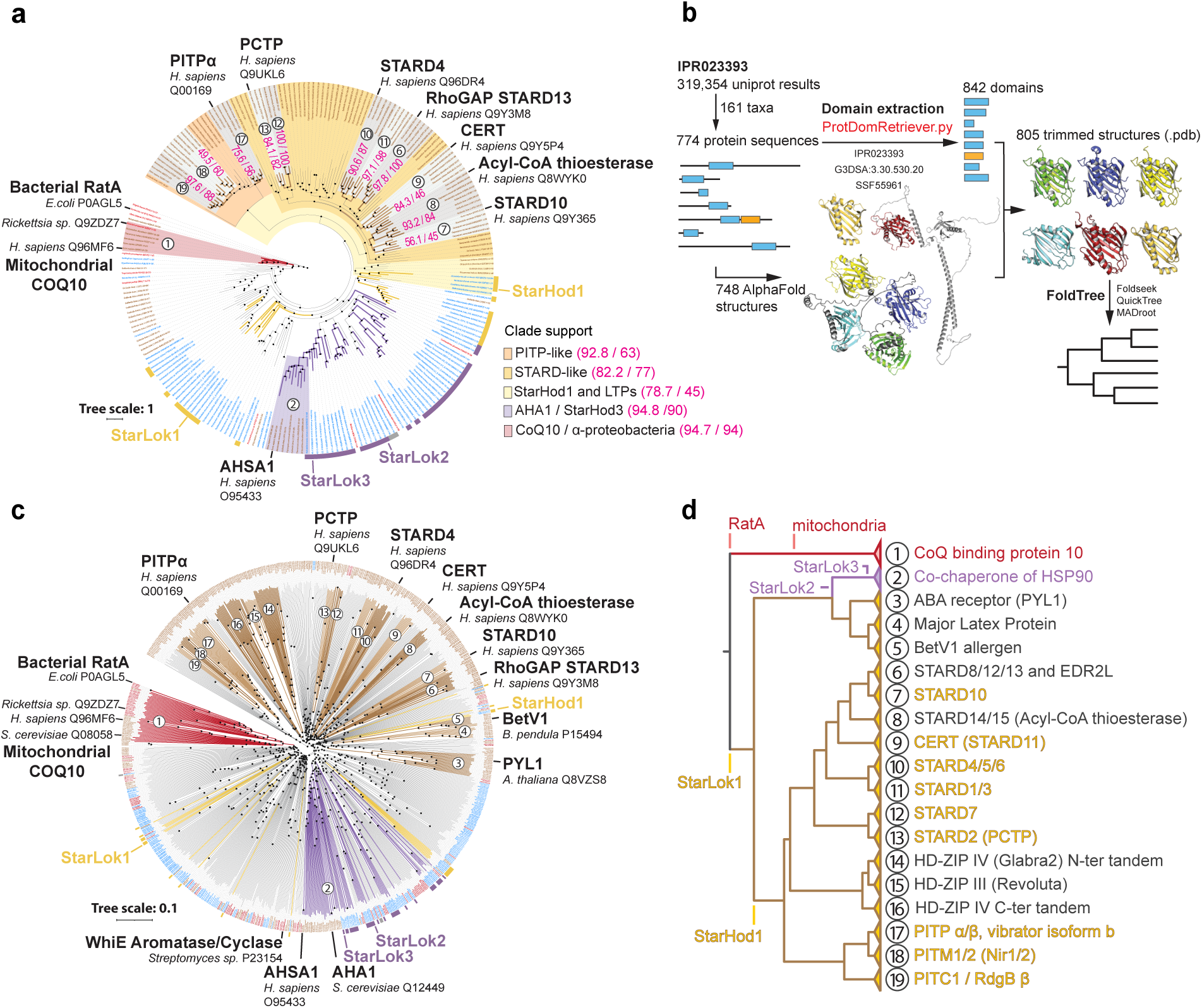
Sequence and structural domain phylogeny support orthology between StarAsg proteins and eukaryotic START domains. **a.** ML phylogenetic tree inferred from 227 START domain sequences, including StarAsg proteins (see Fig. 2a), representative eukaryotes (*Homo sapiens, Nematostella vectensis*, *Capsaspora owczarzaki*, *Dictyostelium discoideum*, *Thecamonas trahens*, *Guillardia theta*), 11 archaeal species, and 25 bacterial species. Sequences were aligned using MAFFT (L-INS-i; --localpair --maxiterate 1,000) with BLOSUM80 and modified gap parameters (--op 1.53 --ep 0.5 --bl 80 --lop −1.0 --lexp −0.5). The best-fit substitution model was selected using ModelFinder according to BIC (Q.PFAM+FO+R5), and ML inference was performed with IQ-TREE. Branch support values correspond to SH-aLRT (%) and ultrafast bootstrap (%) from 1,000 replicates each and are shown in magenta for major clades only. Branch lengths represent substitutions per site. The tree is rooted using *Shigella flexneri* and *Escherichia coli*. Branch-tip labels are colored according to taxonomic origin (brown, eukaryotes; blue, archaea; red, bacteria). Yellow, purple, and gray stripes adjacent to branch-tip labels indicate Asgard sequences belonging to the StarAsg1, StarAsg2/3, or undefined clades, respectively (see Fig. 2a). Red branches highlight the monophyly of mitochondrial CoQ-binding protein 10 homologs with bacterial RatA. Purple, yellow, and brown branches denote StarAsg2/3 and AHA1, StarAsg1, and eukaryotic START LTP clades, respectively. **b.** Workflow for large-scale structural domain phylogeny. ProtDomRetriever was used to identify START domain boundaries using InterPro superfamily accessions, CATH/gene3D annotations, and SUPERFAMILY HMM library accessions. Corresponding structural domains were extracted from AlphaFold models and phylogenetic inference was performed using the FoldTree pipeline. **c.** Rooted structural phylogenetic tree of 805 START domains from 161 species. Visual annotations (branch colors, label colors, and stripe coding) follow the scheme described in panel a. Each eukaryotic START domain class is numbered 1-19 according to the protein-family and class labels. Nodes are represented as black dots. The scale bar indicates substitutions per site. The number of species in each clade is indicated in parentheses. **d.** Simplified cladogram derived from panel c. LTPs previously characterized *in vitro* are highlighted in yellow, signifying the major eukaryotic START domain protein families. Monophyletic classes are labeled using the same numbering as in panel c and named according to established protein-family and class annotations.

While conclusions were consistent across both sequence and structural phylogenetic approaches, FoldTree analysis resolved the evolutionary history of the START domain with deeper resolution and accurately recovered known taxonomic relationships (**Supplementary Figs. 13, 14**). This is likely due to the high sequence plasticity in START domains – less than 31% of proteins passed the χ^2^ composition test – that resulted in high-entropy and gap-rich sequence alignments. However, the overall protein fold remains constant in this superfamily, making it well-suited for structural phylogenetics. Importantly, FoldTree analysis resolved the eukaryotic diversification of START LTPs into previously described subclasses, whose topology is congruent with accepted species relationships (**Supplementary Figs. 13, 14**). These relationships are further described in the **Supplementary Text**. Several of these previously established classifications receive low branch support in ML trees, consistent with the challenging nature of START domains for sequence phylogenetics.

All our trees estimated an ancient origin of the START domain that preceded eukaryogenesis. Sequence alignment-based ML phylogeny (**Fig. 5a**) and structure-based trees (**Fig. 5c**) summarized by a pruned cladogram (**Fig. 5d**) showed a distinct branching group consisting of mitochondrial Coenzyme Q-binding proteins COQ10 (COQ10-like class) and bacterial Ribosome association toxin A (RatA), consistent with previous groupings^62^. The divergence of COQ10 from alphaproteobacterial RatA homologs was also well supported (SH-aLRT/UFBoot = 94.7/94) in ML trees (**Fig. 5a**). These analyses reflect a likely bacterial origin of the nucleus-encoded, mitochondrially localized COQ10 protein.

Our analyses also supported archaeal origins for the Hsp90 co-chaperones AHA1 (AHSA1/AHSA2 in human and Aha1 in yeast). Phylogenetic analyses indicate that AHA1-like proteins originate within Asgards and form a monophyletic group including nine StarAsg3 homologs. The grouping of AHA1 and StarHod3 is well supported (SH-aLRT/UFBoot = 94.8/90) and this clade is within the broader StarAsg3 lineage with lower support (93.8/79) (**Fig. 5a**). In the full dataset, StarAsg3 and AHA1 form a monophyletic group with StarAsg2-like proteins, albeit with low support (**Fig. 5a**, **Supplementary Fig. 6**). In contrast, when restricting the analysis to prokaryotic START domain only, StarAsg2 and StarAsg3 form a strongly-supported monophyletic clade (97.5/92; **Supplementary Fig. 11**), consistent with earlier analyses (**Extended Data Fig. 2**). Thus, AHA1-like proteins are likely to have been inherited from an ancestral StarAsg3 lineage in Asgards.

In eukaryotes, START domains are the basis for multiple classes of LTPs that shuttle lipids intracellularly. Although the available phylogenetic signal did not allow a definitive assignment of START domain LTPs to an Asgard origin, both the sequence-based reconstructions and the FoldTree analysis consistently positioned Heimdallarchaeia StarAsg1-like sequences near the base of eukaryotic LTP diversity. All the LTP START domains in our dataset root with four such sequences (StarHod1/A0A1Q9NB21, A0A1Q9P3Z5, A0A1Q9NZ53 and A0A1Q9NLH4), which themselves form a monophyletic group associated with eukaryotic START LTPs (**Fig. 5a, 5c**). In alternative reconstructions, individual StarAsg1 sequences clustered within the eukaryotic LTP assemblages (**Supplementary Fig. 8**). When restricting the analysis to eukaryotic START LTPs and StarAsg1 homologs (**Supplementary Fig. 7**), StarAsg1 sequences formed a supported clade (97.9/89) together with Bet v 1-like proteins, including plant PYL1; this combined assemblage connects to the broader STARD-like LTP and PITP radiation, but with only weak support (58.5/53). All StarAsg1-like sequences belong to a group characterized by an enlarged predicted binding cavity, a structural feature that is further enhanced in eukaryotic START LTPs (**Extended Data Fig. 3a**). Structurally related START proteins were identified in other archaeal lineages (**Fig. 2d**), several of which grouped with StarAsg proteins in Foldseek-based structural analyses (**Extended Data Fig. 3b**). However, these non-Asgard proteins display smaller or poorly defined predicted binding cavities (**Extended Data Fig. 3c-d**), suggesting that they may not function as LTPs, though fully defining the proteins requires additional analyses.

Based on these analyses and the established relationship of eukaryotes within Asgard archaea, the most parsimonious interpretation is that eukaryotic START domain LTPs could have diversified from an inherited StarAsg1 protein (**Extended Data Fig. 4**). Derivation of eukaryotic LTPs from StarAsg2/3 (0/51; **Supplementary Fig. 9**) or from bacterial START domains (**Supplementary Fig. 10**) was not supported, whereas the grouping of StarAsg1 with eukaryotic LTPs was consistently recovered across ML reconstructions despite weak branch support (**Supplementary Figs. 7, 8**), as well as in the FoldTree analysis. In this potential topology, the smaller-cavity StarAsg2/3 proteins diverge from within the enlarged-cavity assemblage (**Figs. 5a, 5c)**, consistent with the cavity-size distribution (**Fig. 2b, Extended Data Fig. 3a**) and implying a loss rather than gain of the lipid-binding cavity.

## Discussion

Lokiarchaeia, and many other Asgards, contain multiple proteins with START domains, the same fold that forms the basis for a large group of eukaryotic LTPs. One of these families, StarAsg1, binds phospholipids and transports them between membranes. Similarities between StarAsg1 proteins and START domain-containing eukaryotic LTPs suggest that the latter could have emerged from protein(s) within the StarAsg1 class inherited during eukaryogenesis. Due to limited branch support in sequence-based phylogenies, key evidence for this relationship was derived from experimental data and structure-based trees. A second class of Asgard START domains, StarAsg3, includes the ancestral proteins for AHAs, essential co-chaperones in eukaryotes. A third class of START domain proteins conserved in eukaryotic mitochondria was inherited from their alphaproteobacterial ancestor^62,63^. To specialize for function as an LTP, START domains would need to adopt enlarged hydrophobic binding cavities lined by a membrane-interacting motif, and that is the feature we observe in StarAsg1 proteins compared to StarAsg2/3. More broadly, an initial analysis based on predicted structures indicates that an increase in binding cavity size is a general feature for the specialization of START domains for lipid binding across both Asgards and eukaryotes (**Extended Data Fig. 3a**).

Bound ligands and transported substrates of StarLok1 are phospholipids with a range of polar headgroups. One of these, PS, has a clear archaeal equivalent in archaetidylserine^64^. Lokiarchaeon MK-D1 contains a phosphatidylserine synthases homolog (A0A5B9DD23) containing the key the residues for L-serine binding (**Extended Data Fig. 5**) and thus is likely to synthesize PS-like phospholipids. These could thus represent native substrates for StarLok1. StarAsg1 proteins also bind the yeast PM, which is enriched in PS and other anionic phospholipids. The ubiquitous polarity of this protein, containing basic patches near its ligand-binding pocket, suggests that an anionic inner PM is widespread in Asgards. Notably, an anionic cytosolic leaflet coupled to a neutral exoplasmic leaflet is a hallmark of the eukaryotic PM and is essential for phagocytosis and vesiculation^65–68^. It is possible that this feature has its origins in archaea, given the noted asymmetric nature of bolalipids that contain anionic phosphoheadgroups on the cytosolic leaflet and neutral glycosylations on exoplasmic leaflets^69^. In this model, eukaryotic cytosolic proteins that interact with membranes, like LTPs, might have originated from the Asgard archaeal ancestor, much like other ESPs involved in membrane remodeling.

We propose that the first eukaryotic LTPs could have acted as metabolic facilitators by fortuitously transferring newly synthesized lipids between membranes, enabling the proliferation of intracytoplasmic compartments. Ancestral START LTPs could have fulfilled this function with their large, hydrophobic binding cavity and membrane-binding capacity. Later, the emergence of more sophisticated LTPs, such as the Osh/ORP proteins that counter-exchange lipid ligands for PI(4)P^70^, would have enabled the generation of organelles with specialized lipid compositions. Bridge-like LTPs, such as those in the VPS13 class, could have emerged due to the need for rapid membrane proliferation in emerging aspects of eukaryotic cell biology, including mitochondrial network extension^71^, the generation of autophagosomes^72^ or spores^73^, and were later adapted to support routine lipid transfer^74,75^ in larger, increasingly compartmentalized cells. In this way, the increasing complexity of eukaryotic cell biology could have been enabled by biochemical advances in the ability of proteins to transport lipids.

## Supporting information

Combined Supplementary Information

## Acknowledgments

Yongxuan Su and the UCSD Molecular Mass Spectrometry Facility aided in analyses. Dennis, Deshmukh, Devaraj, Ghosh and Herzik labs provided instrumentation support. Paula Whelander, Brett Baker, Laura Eme and Buzz Baum provided helpful discussions. The work was supported by the National Institutes of Health (GM142960), the Moore–Simons Project on the Origin of the Eukaryotic Cell (GBMF9734, grant DOI 10.37807/GBMF9734), and the Paul G. Allen Family Foundation.

## Data Availability

Alignments and trees generated are available at iTOL (https://itol.embl.de/shared/BudinLab), scripts used at GitHub (https://github.com/NicoFrL/ProtDomRetriever and https://github.com/NicoFrL/MAFFT_ScoreNGo). Data files used to generate figures will be made available on FigShare.

## Author contributions

N.-F.L. and I.B. conceived and designed the study. N.-F.L. performed experiments, analyzed data, and developed the computational tools. E.K. carried out lipidomic analysis. N.-F.L. wrote the manuscript. N.-F.L. and I.B revised the manuscript. I.B. supervised the project and acquired funding.

---

**Extended Data Fig. 1.**
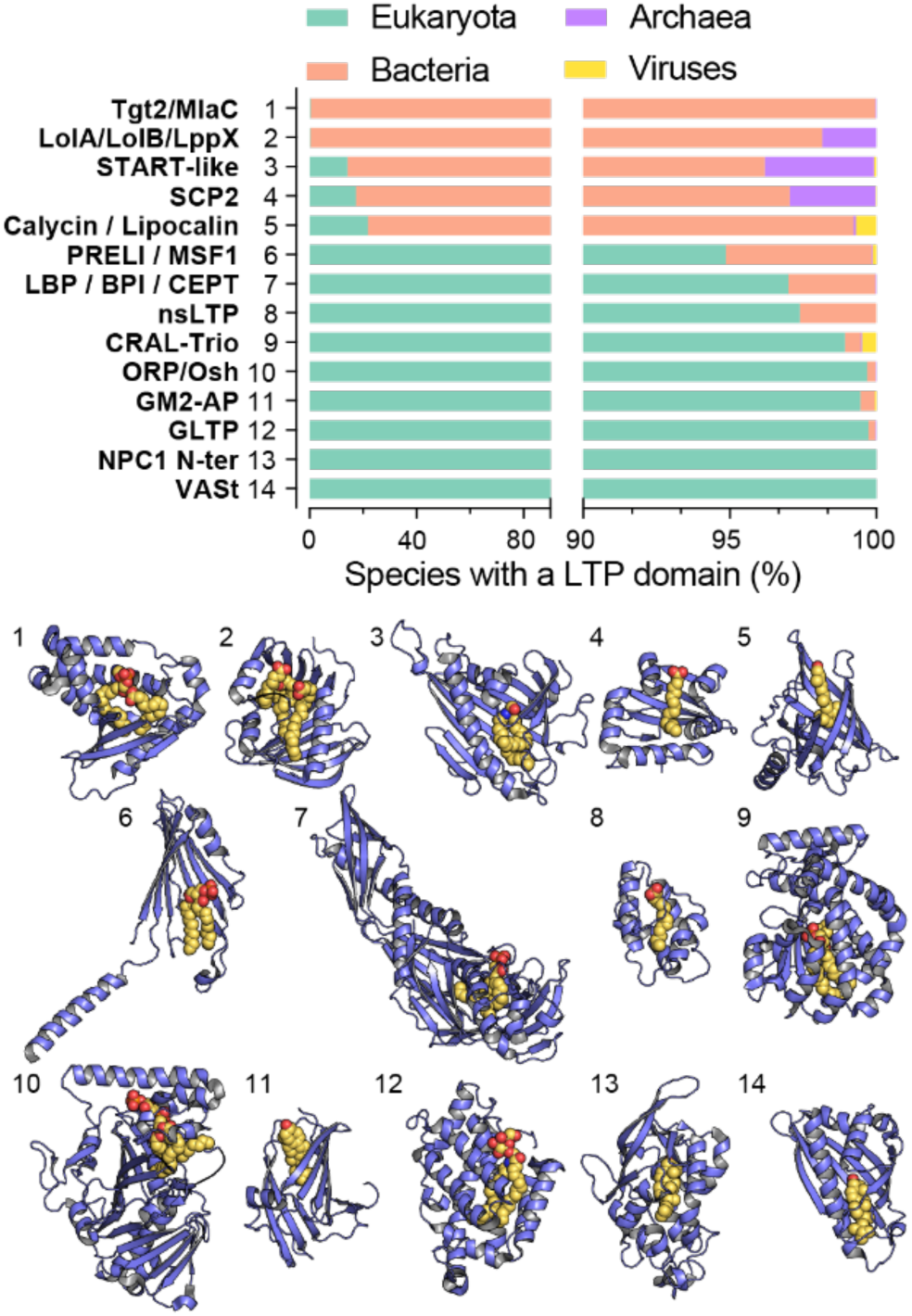
Distribution and structural diversity of shuttle-like LTPs. *Top panel:* The proportion of total sequenced genomes that express shuttle-like LTP classes belonging to each domain of life. Most known LTP classes are exclusive to eukaryotes; archaea contain examples of three (2-4). The data are split into two segments to highlight distributions under 10%. *Bottom panel:* Structures of example proteins from each shuttle-like LTP class shown in the top-panel, containing a bound lipid ligand (alkanes, yellow; oxygen, red; nitrogen, blue; phosphate, orange). 1. MlaC (lipid: phosphatidylethanolamine (PE), pdb: 7VR6). 2. Lppx (α-linolenic acid and docosahexaenoic acid, 2BY0), 3. START domain of CERT (ceramide, 2E3P), 4. SCP-2 (palmitate, 4JGX), 5. RBPA (retinol, 1RBP), 6. Ups1 (phosphatidic acid, 4YTX), 7. BPI (phosphatidylcholine (PC), 1BP1), 8. nsLTP (palmitoleate, 1FK3), 9. Sec14 (PC, 3B7Z), 10. Osh4 (phosphatidylinositol 4-phosphate (PI4P), 3SPW), 11. NPC2 (ergosterol, 6R4N), 12. GLTP domain of 4-phosphate adaptor protein 2 (FAPP2) (*N*-oleoyl-galactosylceramide, 5KDI), 13. N-terminal domain of NPC1 (cholesterol, 3GKI), 14. Lam2 (ergosterol, 6CAY).

**Extended Data Fig. 2.**
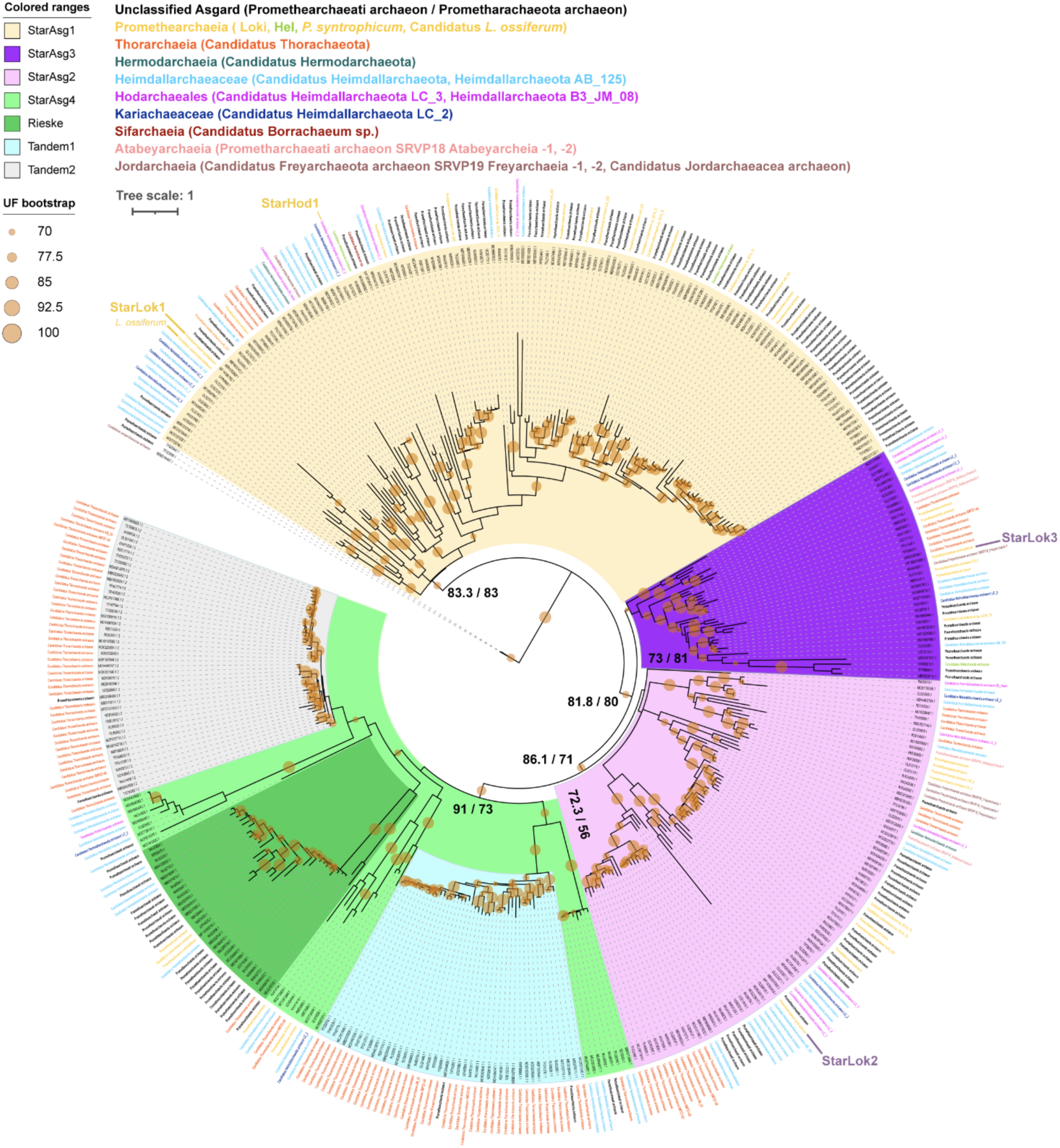
Phylogenetic tree of START domains identified in Asgards. ML tree (IQ-TREE, WAG+F+I+R5) of trimmed START domains in Asgard proteomes. Each color indicates a START domain subclass. Tandem1 and Tandem2 belong to the same protein, and together with Rieske define the class StarAsg4. Branch support is shown as brown circles when both bootstrap and SH-aLRT values are ≥ 70. Relevant support values are reported as SH-aLRT/ UFBoot values respectively. Each species name is colored according to their respective Asgard clade as in the legend. The positions StarLok1/2/3 and StarHod1 are indicated. The StarAsg1 homolog in the isolated strain Candidatus *Lokiarchaeum ossiferum* is indicated as a direct sister of StarLok1.

**Extended Data Fig. 3.**
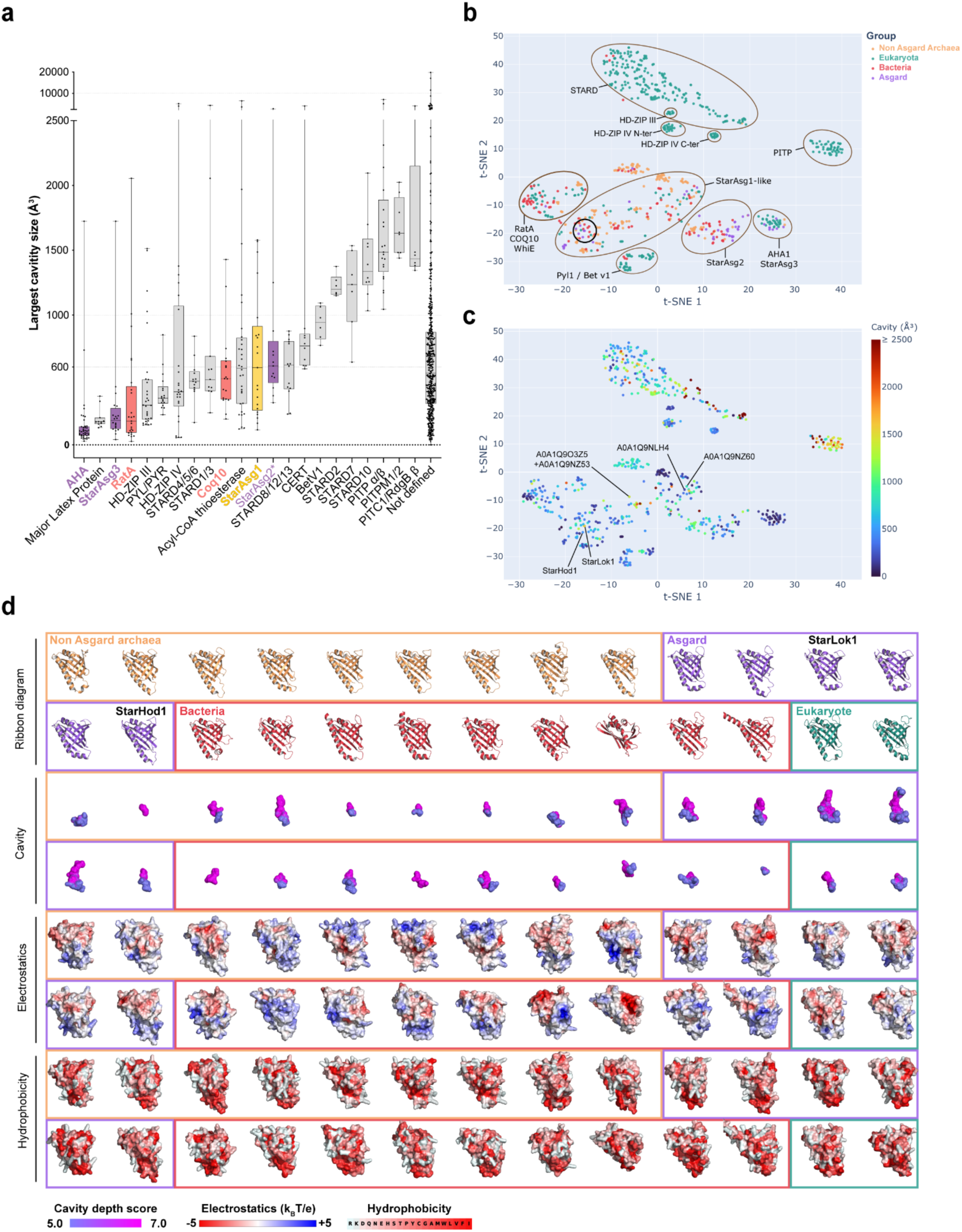
Cavity size distribution of each orthologous group of 805 START domain spanning the tree of life. **a.** Each dot represents the largest cavity size identified in each START domain. Data are presented as box plots, showing the minimum, maximum, median, 25th percentile, and 75th percentile values for cavity sizes within each phylogenetic class. **b.** t-SNE projection of 805 START-domain protein structures based on all-versus-all structural similarity (Foldseek TM-scores), colored by taxonomic group. Proteins with similar three-dimensional folds cluster together (brown ovals), revealing distinct structural families annotated based on known homologs. The black circle indicates a protein cluster within 2.5 t-SNE units of StarLok1 (n=26), shown in detail in panel **d**. **c.** Same t-SNE projection as in **b**, colored by ligand-binding cavity volume (Å³, capped at 2500 Å³ for visualization). StarLok1, StarHod1 and other StarAsg proteins phylogenetically close to eukaryotic LTPs are indicated. STARD-domain proteins cluster in the upper region and display a broad range of cavity sizes, while several Asgard-specific proteins (StarAsg, StarLok1, StarHod1) exhibit cavity volumes comparable to their eukaryotic counterparts. **d.** Structural and physicochemical properties of the 26 START-domain proteins within the t-SNE vicinity of StarLok1. For each protein, ribbon diagrams, ligand-binding cavities (violet to magenta surface, score 5-7), electrostatic potential surfaces (red, negative; blue, positive; −5 to +5 k_B_T/e) and hydrophobicity surfaces (Einsenberg scale) are shown. Proteins are grouped by taxonomic origin: Non-Asgard Archaea (light orange), Asgard (purple), Bacteria (red) and Eukaryota (teal). The Asgard proteins all feature enlarged, well-defined binding cavities and polarized electrostatics. In contrast, most non-Asgard archaeal and bacterial proteins within this cluster show small or poorly-defined predicted binding cavities, and increased variability in surface electrostatics relevant for lipid binding and membrane interactions, respectively.

**Extended Data Fig. 4.**
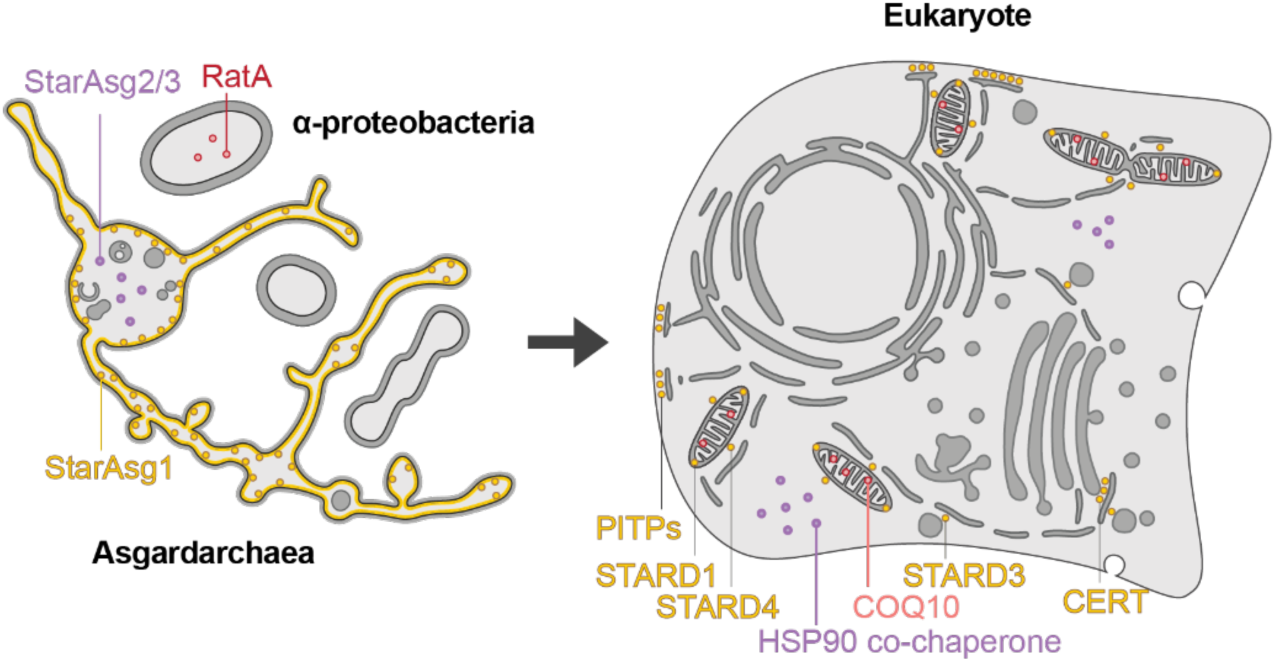
Proposed relationships between START proteins in the prokaryotic ancestors of eukaryotic cells (left) and their localization in eukaryotes. StarAsg1 is a putative membrane-bound lipid-binding protein in Asgards, which could share a common ancestry with several START domain proteins acting as LTPs. These are predominantly localized to membrane contact sites between organelles. StarAsg2 and StarAsg3, proteins of unknown functions in Asgards, share an origin with the conserved HSP90 co-chaperone in the cytosol of eukaryotes. RatA, a START domain protein in alpha-proteobacteria, shares ancestry with the mitochondrial protein COQ10.

**Extended Data Fig. 5.**
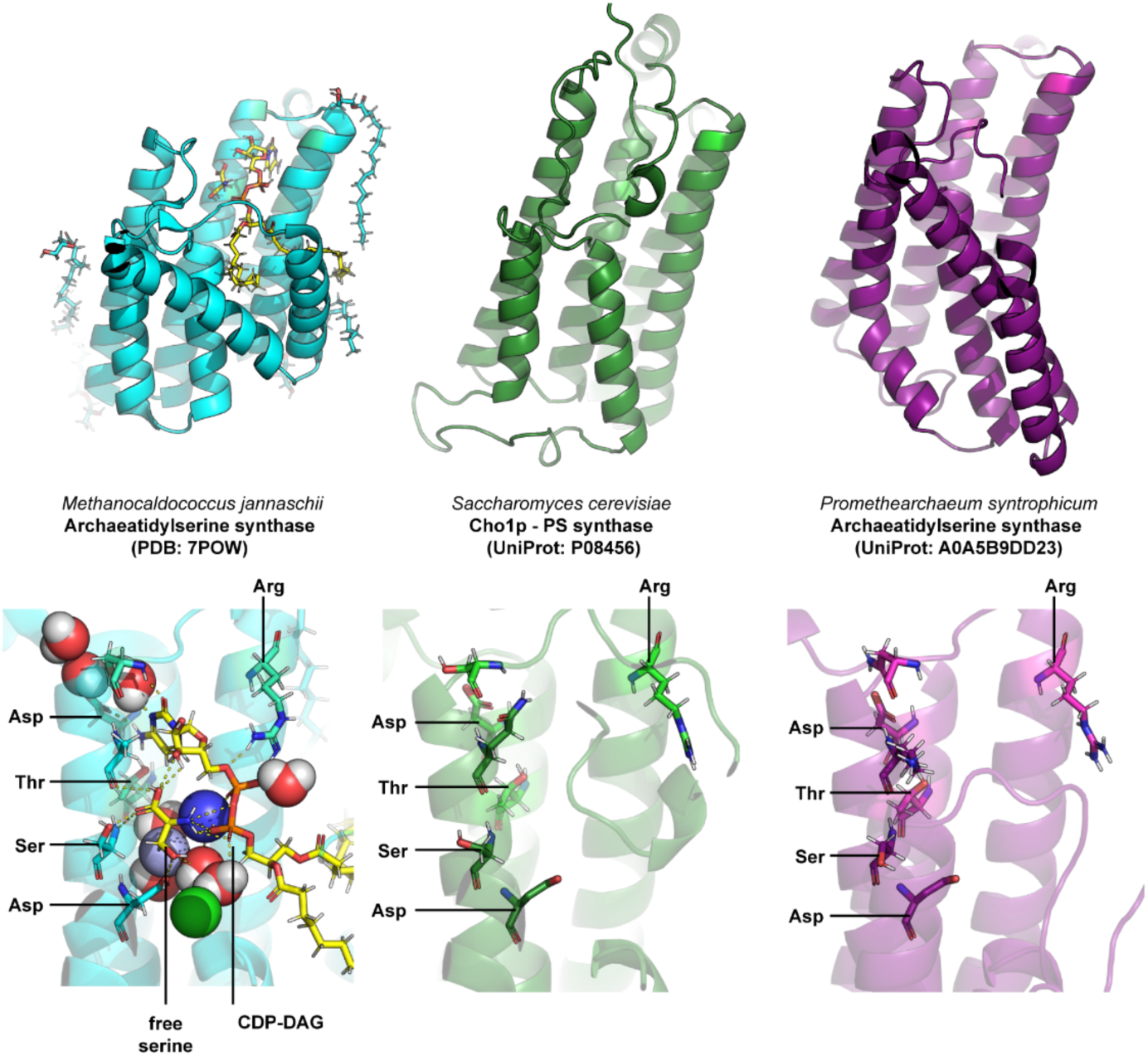
Structural conservation supports serine-dependent anionic lipid synthesis in Asgard archaea. Overall structures of *Methanocaldococcus jannaschii* archaeatidylserine synthase (PDB: 7POW) and AlphaFold models of *Saccharomyces cerevisiae* Cho1p phosphatidylserine synthase (UniProt: P08456) and *Promethearchaeum syntrophicum* archaeatidylserine synthase (UniProt: A0A5B9DD23) are shown in the upper panels. The *M. jannaschii* enzyme is displayed in complex with CDP-diacylglycerol (CDP-DAG) and free L-serine, illustrating substrate positioning within the catalytic pocket. Close-up views highlight conserved residues involved in catalysis and substrate coordination (Arg, Asp, Thr, and Ser). Equivalent residues are preserved in the Asgard homolog, where the spatial arrangement of catalytic side chains closely mirrors that observed in the yeast phosphatidylserine synthase. The conservation of active-site architecture supports the hypothesis that Asgard archaeal enzymes may catalyze serine-dependent synthesis of anionic phospholipids, consistent with the potential presence of phosphatidylserine-like lipids in Asgard membranes. Water molecules (oxygen in red, hydrogen in white), calcium ions (dark blue), magnesium ions (light blue), and chloride ions (green) are shown as spheres. Amino acid residues of interest are shown as sticks, with oxygen atoms in red and nitrogen atoms in blue.

## Methods

### Nomenclature

START: StAR-related transfer, it refers to START-like domain superfamily *stricto sensu (IPR:* IPR023393). In opposition to other “Starkin” domains such as members of VAD1 Analog of StAR-related lipid transfer family and PRELI/MSF1, both display a helix-grip fold analogous to START domains. StAR: Steroidogenic acute regulatory protein. SRPBCC: NCBI Conserved Protein Domain superfamily associated with the START domain (NCBI: cl14643), START/RHO_alpha_C/PITP/Bet_v1/CoxG/CalC ligand-binding domain superfamily. STARD: (StAR)-related lipid transfer domain.

### Lipid transfer protein classification analysis

The 14 classes of LTP we identified were previously described^76–80^. The Interpro accession number of each of these classes was used to determine the number of taxonomic species expressing at least one of these proteins: 1) Tgt2/MlaC (Superfamily IPR042245), 2) LolA/LolB/Lppx (Superfamily IPR029046), 3) START-like domain (Superfamily IPR023393), 4) SCP2 sterol-binding domain (superfamily IPR036527), 5) Calycin-Lipocalin (superfamily IPR012674), 6) PRELI/MSF1 (Domain IPR006797), 7) LBP/BPI/CEPT (i.e. Bactericidal permeability-increasing protein, alpha/beta domain; Superfamily IPR017943), 8) nsLTP (i.e. Bifunctional inhibitor/plant lipid transfer protein/seed storage helical domain; Superfamily IPR036312); 9) CRAL/Trio (Superfamily IPR036865), 10) ORP/Osh (Superfamily IPR037239),11) GM2-AP (Superfamily IPR036846), 12) GLTP (Superfamily IPR036497), 13) NPC1 N-ter (i.e. Niemann-Pick C1, N-terminal; Domain IPR032190), 14) VASt (i.e. VAD1 Analog of StAR-related lipid transfer; Domain IPR031968).

### Expression and purification of StarLok1

A codon-optimized 444 bp cDNA coding for StarLok1 (UniProtKB: A0A5B9D5D6) C-terminally fused to a thrombin cleavage site (LVPRGS) followed by a 6xHis tag was synthesized and cloned into a pET-21a(+) vector (GenScript) and expressed in BL21-Gold(DE3) (Agilent). StarLok1 was purified either from native soluble fraction or refolded from an insoluble fraction. For native purification, a 1 L culture LB Lennox with 50 *µ*g/mL ampicillin was grown to an OD_600_ of 0.5 at 37°C then induced (0.2 mM IPTG) before overnight growth at 16°C. Cell pellets were resuspended in cold TN buffer (50 mM Tris, 500 mM NaCl, pH 8.0, at 4°C) supplemented with protease inhibitor cocktail (Roche cOmplete EDTA-free, 10 µM bestatin, 1 µg/mL pepstatin A, 10 µM phosphoramidon), 10 mM imidazole, 1 mM DTT, 20 µg / mL of DNase I (Roche), 5 mM MgCl_2_ and 1 mg/mL of lysozyme. Cells were then disrupted by sonication and the lysate was clarified by ultracentrifugation (186,000 x *g,* 1h, 4°C). Fifty volumes of supernatant were applied to one volume of Ni-NTA resin slurry (cOmplete His-Tag Purification Resin) and incubated overnight at 4°C. The resin suspension was applied to an Econo-Pac chromatography column (Bio-Rad) successively washed with TN containing 10 mM and 20 mM imidazole. The protein was eluted with TN containing 250 mM imidazole and the eluate concentrated at 2500 *x g* using Amicon Ultra centrifugal unit (MWCO 3000). The protein was further purified by SEC on an Akta pure 25 FPLC with a Sephacryl S200 HR column equilibrated with TN buffer at a flow rate of 1mL/min and fractionation volume of 2.5 mL. Glycerol was added to the buffer (10% v/v) and protein-containing fractions were pooled and concentrated up to 20 *µ*M. Concentration was determined by both A_280_ (DeNovix DS-11 spectrophotometer) and Sypro Orange (ThermoFisher) staining of SDS-PAGE using BSA standards and samples were frozen in liquid N_2_ and stored at −80°C.

For refolding of insoluble protein, StarLok1 expression was performed as described in the previous section with minor modification. When OD_600_ reached 0.6, 1 mM IPTG was added into the culture that was then incubated three hrs at 37°C before collecting the cells and preparing the pellet for freezing. Pellets were resuspended in 50 mM HEPES-KOH pH 7.00 with a protease inhibitor cocktail (Roche cOmplete EDTA-free, 10 µM bestatin, 1 µg/mL pepstatin A, 10 µM phosphoramidon) and passed 4 times through a high-pressure homogenizer at 1,000 bar (EmusiFlex-c3, Avestin). 20 µg/mL of DNase I (Roche), 5 mM MgCl_2_ were added, and the inclusion bodies were then pulled down by ultracentrifugation (186,000 x *g*, 1 hr, 4°C). The supernatant was discarded, and the white pellet was solubilized in HU buffer (50 mM HEPES-KOH pH 7.0 with 6M urea) by mild vortexing and pipetting and one passage through high-pressure homogenizer (1,000 bar). After centrifugation (186,000 x *g,* 1 hr at 4°C), 10 mM imidazole was added to the clear supernatant, which was applied to 50:1 to Ni-NTA resin slurry (cOmplete His-Tag Purification Resin) and incubated 3 hr at 21°C. The suspension was applied to an Econo-Pac chromatography column (Biorad), washed twice with 10 bed volumes of HU with 20 mM imidazole, and eluted with eight bed volumes of HU with 250 mM imidazole. Refolding was performed by dialysis (3.5K MWCO, Slide-A-Lyzer™ MINI Dialysis Devices) twice against 150 volumes of acetate buffer (164 mM acetic acid (OHAc), 36 mM sodium acetate (NaOAc), pH of 4.04) over 14 hr. Screening for refolding conditions was initially done by light microscopy (ThermoFisher EVOS) after low-volume dialysis. Refolded samples were further purified by FPLC on a HiPrep 16/60 Sephacryl S200 HR column (Cytiva) equilibrated with acetate buffer at a flow rate of 1 mL/min and fractionation volume of 2.5 mL. Purification and refolding were monitored by SDS-PAGE. This protocol yielded 5 to 7 mg of protein per 1 L of bacterial culture, as determined by A_280_.

For detergent-solubilized protein used in lipid transfer assays, expression in *E.coli* BL21 was induced with 1 mM IPTG once the OD_600_ reached 0.95 and incubated for 3 hr. Bacterial pellets were collected as previously described. Bacterial pellets were resuspended in PBS with 1% N-lauroyl-sarcosine (NLS) and 10 mM imidazole, 5 mM MgCl2 and 20 µg / mL of DNAse I. Bacteria were lysed by sonication and the lysate was clarified by ultracentrifugation (186,000 x *g*, 1 hr, 4°C). The supernatant was then applied to a Ni-NTA resin slurry and incubated for 3 hr. After two successive washes (PBS, 1% NLS, 10 mM imidazole), StarLok1 was eluted with 250 mM imidazole. The eluate (12 mL) was adjusted with PBS buffer and no detergent, and concentrated to 2 mL using an Amicon Ultra centrifugal unit (MWCO 3000), diluted again to 13 mL with PBS buffer and concentrated once more until the solution started to oil-out around 500 µL. The protein concentration (1.53 mM) was determined by absorbance spectrometry at 280 nm. The sample was stored at 4°C, and purity and stability were determined by SDS-PAGE prior NBD-lipid transfer assay.

### Expression and purification of StarLok2 and StarLok3

A codon optimization 489 bp cDNA coding for StarLok2 (UniProtKB: A0A5B9DET0) and 432 bp coding for StarLok3 (UniProtKB: A0A5B9DAG8), each N-terminally fused to a 6xHis tag followed by a thrombin cleavage site (LVPRGS), were synthesized and cloned into a pET-28a(+) vector (GenScript) and expressed in BL21-Gold(DE3) (Agilent). Each protein was purified from 1 L culture in its native form as described for StarLok1 in TN buffer supplemented with 1 mM DTT. In this standard condition, the protocol yielded more than 10 mg of protein per 1 L of bacterial culture. Protein concentration and purity were controlled by SDS-PAGE stained with InstantBlue (Abcam).

### Circular Dichroism

CD was performed on a Jasco J-1500 spectrometer using a 350 µL synthetic quartz glass cell with a 1 mm path length (CV1Q035AE2, ThorLabs). Measurements were taken of StarLok1 in either 20 mM Tris, 120 mM NaF, pH 7.4 (native), or in a solution prepared by diluting the stock refolded protein (HOAc/NaOAc, pH 4.0) in milli-Q water. Each spectrum represents the average of five continuous scans recorded from 185 to 260 nm with the following parameters: bandwidth of 1 nm, step size of 0.1 nm, scan speed of 50 nm·min−1, CD scale of 200 mdeg/0.1 dOD, and digital integration time (D.I.T) of 4 s. A control spectrum of buffer samples without protein was subtracted from each protein spectrum. For temperature interval scans, an equilibration time of 20 s was set before each scan at 10°C intervals. The measured ellipticity was converted to molar ellipticity, considering the cuvette path length and the mean residue concentration of proteins. For secondary structure determination, the spectra were analyzed in the 185–240 nm range using the SELCON3 algorithm and the SP175t dataset on the DICHROWEB server^81^ and compared with the algorithm available on the BeStSel web server^82^. Thermal denaturation results were displayed using matplotlib.

### Protein structure analysis

Structures were rendered using PyMOL (v2.5.8, Schrödinger, LLC). Structure alignment comparisons were conducted using the cealign algorithm and the RMSD values were reported. Model-template alignments were obtained using SwissModel^83^. Cavities point clouds were calculated using the CavitOmiX plugin (v. 1.0, 2022, Innophore GmbH)^84^ and cavities were calculated using a modified LIGSITE algorithm^85^. To compute the cavity size, a script calling CavFind and LigSite modules from CavitOmiX was used to extract the number of cavities, their respective volumes and ligsite score. Protein electrostatic surface potential was determined using APBS (pdb2pqr)^86^ with a grid spacing of 0.26. Hydrophobicity was determined using the Eisenberg scale^87^. The Positioning of Protein on Membrane (PPM) was determined using PPM 3.0 web server^88^, using a flat theoretical archaeal plasma membrane composed of Phosphatidylglycerol, Cardiolipin, Archaeol and Menaquinone 8 at a ratio 64:7:8:21. Changing the membrane composition did not significantly change the obtained results. The AlphaFold2 structure of StarLok1 with a thrombin cleavage linked to 6x(His)-tag was computed using ColabFold v1.5.5 notebook. The secondary structure composition was determined using both STRIDE^89^ and DSSP algorithm^90^. The theoretical CD spectra of StarLok1 were determined using either SESCA^91^ and PDB2CD^92^ for comparison with experimental measurement.

### Protein mass spectrometry

StarLok1 sample purified in native condition was dialyzed three times (1 hr, overnight, and 1 hr) against 1 L of TN buffer (10 mM Tris-HCl, 50 mM NaCl, pH 7.4) using a 0.5 mL Slide-A-Lyzer cassette, to ensure complete glycerol removal. For StarLok1 purified in acetate buffer, no dialysis was performed. Protein samples were analyzed for purity and mass identification by liquid chromatography with an Agilent 6230 time-of-flight mass spectrometer (TOFMS) with JetStream electrospray ionization source ESI (LC-ESI-TOFMS)

### Liposome preparation

Stock solutions of 1,2-dioleoyl-sn-glycero-3-phosphocholine (DOPC), 1-palmitoyl-2-{12-[(7-nitro-2-1,3-benzoxadiazol-4-yl)amino]dodecanoyl}-sn-glycero-3-phosphoserine (16:0-12:0 NBD-PS), 1-palmitoyl-2-{12-[(7-nitro-2-1,3-benzoxadiazol-4-yl)amino]dodecanoyl}-sn-glycero-3-phosphoethanolamine (16:0-12:0 NBD-PE), 1-palmitoyl-2-{12-[(7-nitro-2-1,3-benzoxadiazol-4-yl)amino]dodecanoyl}-sn-glycero-3-phosphocholine (16:0-12:0 NBD-PC), N-[12-[(7-nitro-2-1,3-benzoxadiazol-4-yl)amino]dodecanoyl]-D-erythro-sphingosine (C12-NBD-Ceramide) and 1,2-dipalmitoyl-sn-glycero-3-phosphoethanolamine-N-(lissamine rhodamine B sulfonyl) (Rhodamine-PE), 1-palmitoyl-2-oleoyl-sn-glycero-3-phospho-L-serine (POPS) in chloroform (Avanti Polar Lipids) were mixed at the desired molar ratio. The solvent was evaporated under vacuum (54.9 mbar) for 30 min on a rotary evaporator at 37°C. Residual solvent was eliminated by further drying in a vacuum chamber for at least 1 hr. The resulting lipid films were hydrated with 2 mL HK buffer (50 mM HEPES-KOH, 120 mM K-acetate, pH 7.4) and vigorously vortexed with glass beads. The resulting multilamellar vesicles suspension was subjected to five freeze-thaw cycles (liquid nitrogen/37°C) followed by extrusion through a 0.2 µm polycarbonate filter using a mini-extruder (Avanti Polar Lipids). Freshly extruded liposomes, prepared on the same day of the experiment and used within two days, were shielded from light were used for all experiments.

### Dynamic Light Scattering

Measurements were performed using a DynaPro instrument with a Flex-99 correlator (ProteinSolutions) and quartz cuvettes. The instrument sensitivity was set to 30% with a maximum acquisition time of 10 s and a signal-to-noise threshold of 2.5. The cuvette holder temperature was maintained at 25°C. The buffer refractive index was set to 1.333. For kinetics experiments, 20 min autocorrelation curves were acquired from 200 nm extruded liposomes (50 μM lipid concentration) in degassed HK buffer (50 mM HEPES-KOH, 120 mM K-acetate, pH 7.4) with 1 μM of StarLok1 in acetate buffer added after 2 min. Alternatively, StarLok2 and StarLok3 were tested. Data filtering and analysis were performed using DYNAMICS software (v.5.25.44) with a maximum Sum Of Square (SOS) limit at 2100, under baseline limit at 0.995, over baseline limit at 1.02, under amplitude limit of 0.05. For the correlation function, the first 2 coefficients were ignored, and the analysis was truncated at channel 120. The autocorrelation functions were then analyzed using the regularization algorithm.

### Lipid transfer assay

NBD-lipid transfer was measured as previously described^59^ with modifications. Fluorescence was recorded using a Cary Eclipse Fluorescence Spectrophotometer (Agilent) equipped with a temperature controller set to 30°C. In a 500 µL quartz SUPRASIL rectangular cuvette (PerkinElmer), stirred with a magnetic bar, 200 µM donor liposomes (L_D_) containing 2 mol% Rhod-PE and 5 mol% NBD-labeled lipid (NBD-PC, NBD-PE, NBD-PS, or NBD-ceramide) were suspended in HK buffer (50 mM Hepes, 120 mM K-acetate, pH 7.4) pre-warmed at 30°C. NBD fluorescence was monitored at 538 nm (emission slit = 5 nm) with excitation at 473 nm (slit = 2.5 nm), photomultiplier voltage of 800 V, and acquisition interval of 0.5 s. At t = 1 min, 200 µM acceptor liposomes (L_A_; DOPC only) were added. After 3 min of slow spontaneous transfer, StarLok1 (1.60 µL; 5 µM final concentration) was injected, resulting in a significant increase in fluorescence indicative of NBD-lipid transfer from L_D_ to L_A_. As a negative control, 1.73 µL of proteinase K-digested StarLok1 was tested. Digested samples were prepared by incubating StarLok1 with Proteinase K (20 mg/mL, RNA grade; Invitrogen) at a 1:12 (v/v) ratio for 1 h at 42°C, followed by heat inactivation for 5 min at 98°C. Complete proteolysis was confirmed by SDS-PAGE analysis. The amount of NBD-lipid transferred was quantified by measuring the loss of FRET between NBD and Rhod-PE, normalized as:

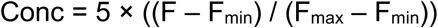

where F is the recorded fluorescence, F_max_ is the mean fluorescence over 30 s prior to protein injection, and F_min_ is the mean fluorescence at equilibrium (t = 12 min, averaged over 3 min) determined in an independent experiment without protein using pre-equilibrated liposome populations L_D_ (2 mol% Rhod-PE) and L_A_, both containing 2.5 mol% NBD-lipid. The coefficient, 5 reflects the 5 µM NBD-lipid accessible in the external leaflet of L_A_. Baseline correction was performed by subtracting the mean normalized signal from control experiments (n = 3) in which 1.64 µL of vehicle buffer (PBS containing 1% N-lauroylsarcosine) was injected instead of protein; no increase in lipid transfer was observed under these conditions. Initial transfer rates (Vᵢ ± SEM) were determined from the slope of a linear fit (least-squares regression, unweighted) to data collected over 3 s post-injection, treating each replicate as an individual data point. Statistical analysis was performed using two-way ANOVA with Tukey’s post-hoc test. All analyses were conducted in GraphPad Prism (v10.2.3).

### Yeast expression

*E. coli* codon optimized sequences for StarLok1, StarLok2, StarLok3 and StarHod1 were cloned into the yeast integration vector pCD256 under a TEF1 promoter and fused with a C-terminal yeGFP. The expression construct was integrated into the *URA3* locus of background strain W303a after double digestion of pCD256 with Pme1/Pac1 and chemical transformation. Clones were isolated by selective plating with G418 and grown in CSM medium with 2.0% w/v glucose. The C2 domain of Bovine Lactadherin N-terminally fused to mRFP was gifted from Sergio Grinstein (Addgene plasmid #74061), and was cloned into a pRS415 vector under the control of a PGK1 promoter and terminator sequences. Yeast cells expressing StarLok1 were transformed and selected on CSM-Leu with 250 µg/mL G418, then grown in CSM-Leu medium with 2.0% w/v glucose. For imaging, stationary phase cells were added to concanavalin-1A-coated 8-well coverslip dishes and imaged with a Zeiss LSM 880 confocal laser scanning microscope equipped with an AxioObserver stand and a Plan-Apochromat 63×/1.4 NA Oil DIC M27 objective. Images were acquired with the following settings: GFP was excited using a 488 nm argon laser at 2.3% power, and fluorescence emission was collected in the 494.95-565.07 nm range using a photomultiplier tube (PMT) detector set at 750 V. Brightfield (BF) images were simultaneously captured using a non-descanned transmission PMT detector. Imaging parameters included a pixel dwell time of 4.1 μs, 8× line averaging, and a confocal pinhole of 0.997 Airy units (50.4 μm) to optimize the signal-to-noise ratio while maintaining optical sectioning. Images were digitized as 16-bit single optical sections with a frame size of 1024×1024 pixels and subsequently cropped to 512×512 pixels for further processing. The physical pixel size was 65.9 nm, yielding a field of view of 33.7×33.7 μm. For super-resolution imaging, the same microscope was used with its Airyscan detector. Z-stacks were collected with a step size of 0.18 μm, covering a total depth of 11.88 μm (66 slices). For optimal resolution, the pixel size was set to 0.04×0.04 μm in the x-y dimension, yielding voxel dimensions of 0.04×0.04×0.18 μm. For dual-channel imaging (GFP/mRFP), Z-stacks were collected with 0.185 μm steps (75 slices, 13.87 μm total depth) at 0.0488×0.0488 μm pixel size, yielding voxel dimensions of 0.0488×0.0488×0.185 μm. Excitation was performed using 488 nm (argon laser) and 561 nm (DPSS laser) at 2% power each. Raw Airyscan images were processed using the ZEISS ZEN software with the standard Airyscan processing algorithm in 3D mode. In FIJI (imageJ), the "Sharpen" filter was applied once to enhance structural details. No additional processing was performed on individual slices to maintain the integrity of the original signal distribution. For visualization purposes, all images were cropped to 512×512 pixels to focus on the regions of interest.

### Isolation of GFP-tagged Asgard archaeal proteins for lipidomic analysis

StarLok1, StarLok2, StarLok3, and StarHod1 GFP-fusion proteins were expressed in S. cerevisiae W303 grown in 50 mL YPD medium at 30°C for 24 h, following 1:100 inoculation from an overnight culture. Cells were harvested by centrifugation (3,000 ×g, 5 min, 4°C), washed once with ice-cold sterile water, and resuspended in 500 µL lysis buffer (10 mM Tris-HCl, 150 mM KCl, 1 mM MgCl₂, pH 7.2; filtered and degassed) supplemented with 1 mM DTT, Complete EDTA-free protease inhibitor cocktail (1 tablet per 20 mL; Roche), and PhosSTOP phosphatase inhibitor (2 tablets per 20 mL; Roche). Cell lysis was performed by cryogenic grinding. The cell slurry was transferred to a ceramic mortar pre-chilled and maintained at −196°C with liquid nitrogen and ground to a fine powder with a pestle. The frozen powder was transferred to a 2 mL microcentrifuge tube and thawed on ice. To reduce viscosity, samples were supplemented with 50 µL DNase I solution (20 µg/mL DNase I Grade II from bovine pancreas [Roche] in 5 mM MgCl₂). Lysates were then clarified by centrifugation (20,000 ×g, 15 min, 4°C), yielding approximately 1 mL of supernatant. For affinity purification, clarified lysate was transferred to a 2 mL round-bottom tube containing 50 µL GFP-Trap Magnetic Particles (M-270; Chromotek), pre-washed and equilibrated in a wash buffer (10 mM Tris-HCl, 150 mM KCl, 1 mM MgCl₂, pH 7.2; filtered and degassed) according to the manufacturer’s instructions. The suspension was incubated for 1 h at 4°C with end-over-end rotation. Beads were washed five times with the wash buffer and stored at 4°C until lipid extraction.

### Analysis of lipids bound to GFP-StarAsgard proteins

Total lipids bound to GFP-StarAsg protein were extracted via an adapted BUME method^93^. Magnetic particles bound to GFP-StarAsgard proteins were resuspended in 500 µL of 3:1 (v/v) butanol:methanol and vortexed until complete resuspension. The suspension was transferred to a 12×75 mm conical glass tube, combined with 500 µL of 3:1 (v/v) heptane:ethyl acetate and 500 µL of 1% acetic acid, and centrifuged for 5 min at 3,100 ×g. The upper phase (450 µL) was transferred to an amber glass vial with a PTFE rubber-lined cap (Wheaton) and evaporated at 40°C for 1 h using a rotary evaporator. The dried lipid extract was resuspended in 50 µL of 18:1:1 (v/v/v) 2-propanol:dichloromethane:methanol and vortexed for 3 min. Samples were centrifuged for 1 min at 3,100 ×g and stored at −20°C until analysis

For lipidomics analysis, deuterated internal standard (Avanti) representing each of the 15 lipid classes analyzed were spiked into the resuspended samples. These allow for semi-quantification of species within these classes. Other potentially present lipid species, such as ceramide derivatives, glycosylated lipids, and non-sterol terpenes, were not quantified using this method. Polar lipids were measured according to a previously described method^94^, with slight modifications. The Vanquish UHPLC (Thermo Fisher Scientific) was interfaced with a QExactive mass spectrometer (Thermo Fisher Scientific). A Waters T3 1.6 µM 2.1 mm x 150 mm column was used for chromatographic separation using a step gradient from 25% buffer A (10 mM ammonium formate and 0.1% formic acid in water) to 100% buffer B (70/30 isopropanol/acetonitrile with 10 mM ammonium formate and 0.1% formic acid) over 35 min. Flow rate was set at 0.3 mL/min. This LC gradient minimizes ion suppression. Lipid samples were analyzed using data-dependent acquisition (DDA) with an NCE of 30 in negative mode and an NCE of 25 in positive mode. Ion source parameters were: Sheath Gas 48 AU; Aux Gas 11 AU; Sweep Gas 1 AU; Spray Voltage 3.5 kV for positive mode and 2.5 kV for negative mode; Capillary Temp. 250°C; S-lens RF level 60 AU; Aux Gas Heater Temp. 413°C. All ions in the mass range of 200-1200 m/z were monitored. MS1 resolution was set at 70,000 (FWHM at m/z 200) with an automatic gain control (AGC) target of 1e6 and Maximum IT of 200 ms. MS2 resolution was set to 17,500 with an AGC target of 5e4, fixed first mass of 80 m/z, and Maximum IT of 50 ms. The isolation width was set at 1.2; Dynamic Exclusion was set at 3 s. Lipids were identified and quantified with Lipid Data Analyzer Software.

### InterPro Sequence alignments and phylogeny of START proteins in Asgards

The initial dataset consists of the 56 identified sequences corresponding to the InterPro accession number IPR023393 in annotated Asgard strains including Lokiarchaeia (MK-D1, CR_4, b31, b32 and GC14_75), Thorarchaeia (SMTZ1_45, SMTZ1_83 and AB25), Heimdallarchaeia (including LC_3 and B3_JM_08 both reclassified as Hodarchaeales, LC_2 as Kariarchaeaceae, and AB_125 as Heimdallarchaeiaceae^25,26^) and Helarchaeales. One sequence (A0A135VUP7) from Thor was removed because it was identical to A0A135VGP2. Full-length protein sequences were aligned using MAFFT L-INS-i with default parameters and Blosum62 matrix^95^. The alignment results were used to compute a ML phylogeny using the PhyML server (PhyML 3.3.2), with Smart Model Selection based on Bayesian Information Criterion (SMS BIC), using a neighbor-joining starting tree (BioNJ)^96^. The inference was then reiterated in IQ-TREE^97^, using the same substitution model, frequency parameters, and seed, and branch support was assessed using SH-aLRT (1,000 replicates and ultrafast bootstrap (UFBoot2, 1,000 replicates). The tree was then displayed and annotated using the iTOL online tool^98^ and a single sequence from Helarchaeales was selected as a root.

### NCBI sequences alignment and phylogeny of START proteins in Asgards

Asgard proteome sequences were retrieved from the NCBI database (GenBank Release 264), comprising 711,493 Identical Protein Groups (IPGs). Protein domain analysis was performed using InterProScan version 5.72-103.0^99^ deployed via Docker Desktop 4.37.2 (179585) on macOS. Analysis was restricted to Gene3D, Pfam, and SUPERFAMILY profile signatures. A custom bash script divided protein sequences into smaller batches and ran analyses in parallel (two concurrent processes, two CPU cores each). The pre-calculated match lookup service was utilized with default configuration. Output files were organized by batch and the 508,901 newly annotated sequences were concatenated for downstream analysis. After assigning taxonomic ranks to each sequence using NCBI annotation, 411 START domains were identified using InterPro annotations. (SSF55961; G3DSA:3.30.530.20; PF10604; PF11485; PF08982; PF06240; PF19569; PF02121; PF08327; PF03364; PF01852; PF00407; IPR023393). Each sequence was trimmed according to the longest non-overlapping domain coordinate given by the InterPro Analysis. A subset of START domain-containing proteins in the Thorarchaeia clade contained two domains; these were retained as distinct protein sequences. The trimmed sequences were aligned using Muscle5^100^. ML phylogenies were inferred with IQ-Tree following model selection using ModelFinder. Branch support was assessed using SH-aLRT (1,000 replicates) and ultrafast bootstrap (UFBoot2, 1,000 replicates). Five monophyletic domain classes were identified. To represent domain abundance, the average number of species in each Asgard archaeal taxon was estimated using seven universal single-copy proteins (SecY, uS13, uS3, uS4, aIF2, Arginine and Serine RNA ligase) among the 508,901 annotated sequences. The mean count of these marker proteins per taxon was used as a proxy for species abundance. START domain abundance in each taxon was normalized by the estimated species abundance.

### Large sequence dataset construction

A set of 82 species of archaea including 14 Asgards, 4 Stygia, 4 Acherontia, 18 Stenoarchaea, 5 Methamonada, 4 Diaforarchaea, 31 TACK group members and 2 DPANN group members, was collected based on recent phylogeny^101^. Similarly, 32 eukaryotes were selected including Obazoa, Amoebozoa, Excavata, Cryptista, Archaeplastida and TSAR reflecting previous molecular-clock analysis^102^. 46 Bacterial species were also selected a posteriori, to increase the depth of the tree by including members of Acidobacteriota, Bdellovibrionota, Campylobacterota, Chrysiogenota, Coprothermobacterota, Deferribacterota, FCB group, Fusobacteriota, Myxococcota, Pseudomonadota (Alpha, Beta and Gammaproteobacteria), PVC group, Terrabacteria group, (Actinomycetota, Deinococcota), Thermodesulfobacteriota and Spirochaetota. For each species, START-containing protein sequences were collected using IPR023393 accession as a request to UniProt and then processed with CD-HIT (-c 0.9 -n 5) to identify putative in-paralog groups^103^. For each of these groups, spurious redundant sequences were manually removed depending on their annotation in genomic databases such as GenBank, Gramene, FlyBase, EnsemblMetazoa and RefSeq. When locus annotations were lacking between every in-paralog, the longest transcript was selected. From this new protein dataset, as a list of accession file, ProtDomRetriever.py was executed using as an input the InterPro entries IPR023393, CATH G3DSA:3.30.530.20, and SUPERFAMILY SSF55961.

### Reduced sequence-based Maximum Likelihood phylogeny of START domains

A curated pool of 842 START domains derived from 774 protein sequences was assembled using ProtDomRetriever.py. From this pool, specific sequence subsets were defined to generate the phylogenetic trees presented in the main and supplementary figures, enabling comparisons across START clades or within major domain groups. The sequences were aligned using MAFFT (v7)^104^. To empirically determine optimal alignment parameters, we developed MAFFT ScoreNGo.py, a wrapper that systematically evaluates alternative MAFFT configurations. The following alignment strategies were tested: G-INS-i, E-INS-i, and L-INS-i, using both BLOSUM62 and BLOSUM80 scoring matrices, together with a range of gap-opening, gap-extension, and local alignment penalty parameters. In total, 337 parameter combinations were evaluated using the standard search mode. Each resulting alignment was scored using a conservation-based metric implementing Jalview’s per-column conservation scheme^105^, derived from the AMAS method^106^. For each alignment column, conservation was evaluated based on agreement across 10 physicochemical property classes, with columns exceeding a defined gap threshold excluded from scoring. Per-column conservation scores (0–11) were summed to obtain a global conservation score for each alignment. For each dataset, the alignment yielding the highest global conservation score was retained for downstream phylogenetic inference. This empirical optimization strategy was designed to maximize residue-property consistency across the conserved START domain core, improving alignment coherence and preserving phylogenetically informative sites. Alternative alignments generated with MUSCLE5 were also evaluated using the conservation-based scoring framework described above. In direct comparisons on selected START domain datasets, MUSCLE5 produced lower global conservation scores and more gap-dense alignments relative to MAFFT under optimized parameters, reducing coherence across conserved core residues. MAFFT was therefore used for phylogenetic inference in the tree-of-life START domain analyses. ML phylogenies were inferred using IQ-TREE with extended model selection implemented via ModelFinder. ModelFinder was used to evaluate a comprehensive set of substitution models, including mixture models (C10, C20, C40, C60), LG4M and LG4X models, alternative equilibrium frequency schemes (F, FO, FQ), nuclear substitution matrices, and multiple rate heterogeneity models (E, I, G, I+G, R, I+R; 4–20 categories). The best-fitting model was selected according to the Bayesian Information Criterion (BIC) for each dataset. Branch support was assessed using SH-aLRT (1,000 replicates) and ultrafast bootstrap (UFBoot2, 1,000 replicates). The trees were displayed and annotated with iTOL^98^.

### Large structural phylogeny of START domains

AlphaFold structures were retrieved from Alphafold Protein Structure database. A trimming algorithm was applied to the PDB structures using the position file generated by ProtDomRetriever, to generate new .pdb files excluding any atoms that are not part of the START domain. When multiple START domains are present in a structure, each of these domains are extracted and copied in an independent new file. This set of trimmed structures was submitted to the FoldTree (build d0f0239) pipeline^60^, using Google Colab Notebook on a Python3 runtime accelerated by a A100 GPU. The rooted tree was then displayed and annotated with iTol without modifying the topology. From this tree, a smaller collapsed cladogram was constructed by deleting all sequences not belonging to a known clade of functional protein. Taxonomic congruence was evaluated by comparing the START domain phylogeny with a synthetic species tree extracted from the Open Tree of Life (OTOL) database^107^ using the induced_subtree API. After pruning to identical taxon sets, topological agreement among major eukaryotic clades and lineage-specific paralogous expansions was assessed using a tanglegram representation.

### Structural comparison analysis

Structural comparisons were performed using Foldseek (exhaustive search mode) to compute all-versus-all TM-scores across 805 trimmed AlphaFold2 structural models. A pairwise distance matrix (1 − TM-score) was constructed and reduced to two dimensions by t-SNE following PCA pre-processing (50 components), using scikit-learn. Ligand-binding cavity volumes were derived from cavity volumes previously computed using CavitOmiX. Figures were generated using Plotly.

