## Supplementary material for "A eukaryotic-like phospholipid transfer protein conserved in Asgard archaea": Combined Supplementary Information

For

This file includes:

- Supplementary Figures 1-14
- Supplementary Text
- References

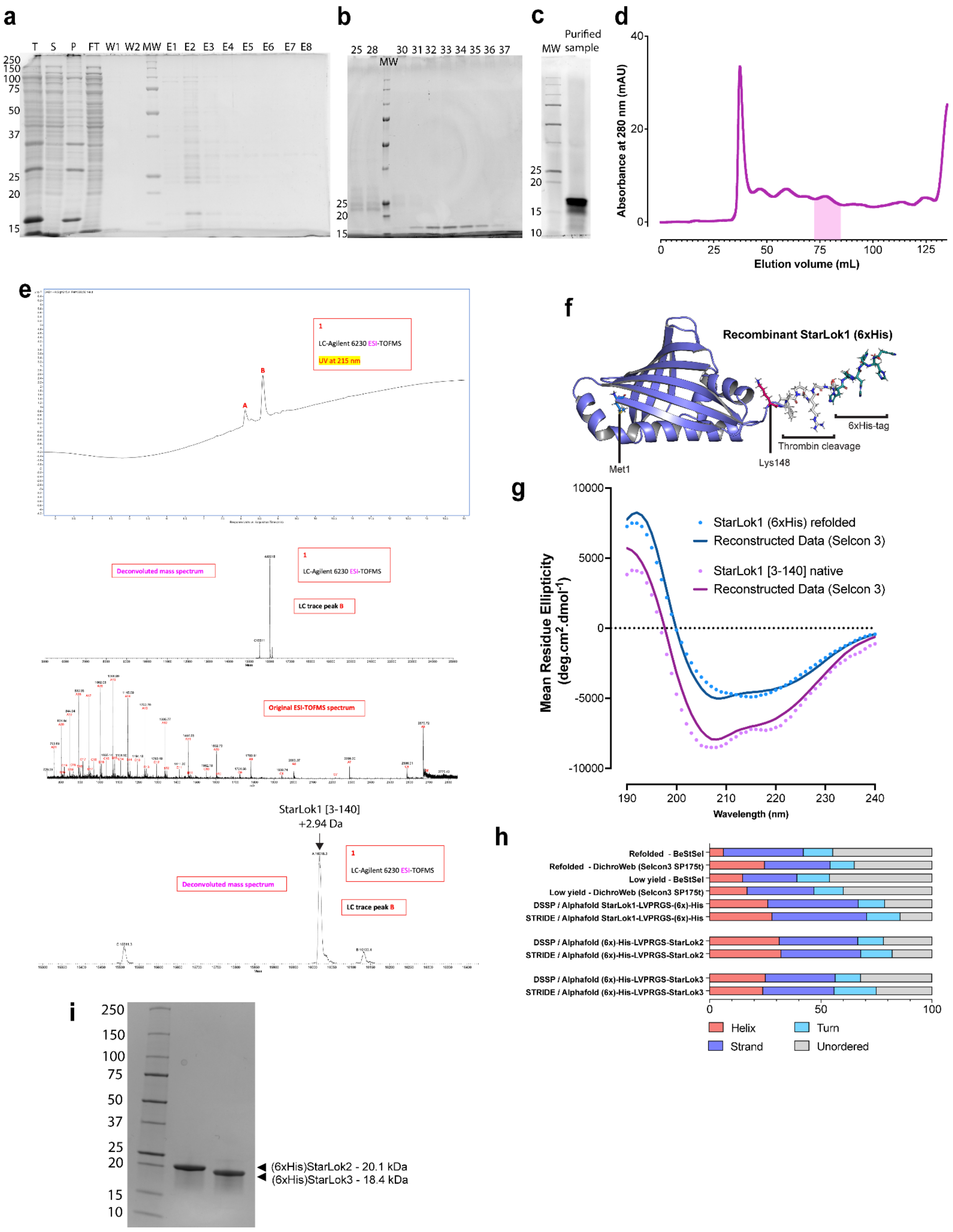


**Supplementary Fig. 1: Purification and folding of StarLok proteins expressed in *E. coli*.**

**a.** SDS-PAGE results of recombinant StarLok1 expression and purification using immobilized metal affinity chromatography (IMAC). Letter indicates purification steps as follows: T: total, S: supernatant, P: pellet, FT: Flow through, W1 and W2: Washes, E1 to E8 corresponds to elution fractions. MW: Molecular weight marker. Numbering on the left indicates the apparent molecular weight in kDa. **b.** SDS-PAGE results from samples recovered from each fraction after size exclusion chromatography. MW: Molecular weight marker. Numbering on the left indicates the apparent molecular weight in kDa. **c.** SDS-PAGE showing 1.5 μL equivalent sample of stock of purified StarLok1. **d.** SEC chromatogram, the plain pink area under the curve indicates the concentrated fraction (31 to 36) containing StarLok1. **e.** LC-ESI-TOFMS analysis of recombinant StarLok1 (6xHis) purified from *E. coli*. The chromatograms depicted the deconvoluted mass spectrum of the main detected peak. **f.** AlphaFold2 model of the recombinant StarLok1 (6xHis). **g.** Circular dichroism spectra of StarLok1 purified natively (blue) and purified at pH 4.0 (pink). Reconstructed data using Selcon 3 algorithms are displayed. **h.** Comparison between the secondary structure determination of StarLok1 obtained from CD spectra with secondary structure distribution of AlphaFold structures of StarLok1, StarLok2 and StarLok3 in the presence of the purification tag predicted by two algorithms (DSSP and STRIDE). **i.** SDS-PAGE showing StarLok2 and StarLok3 (6xHis) purified from *E. coli*.

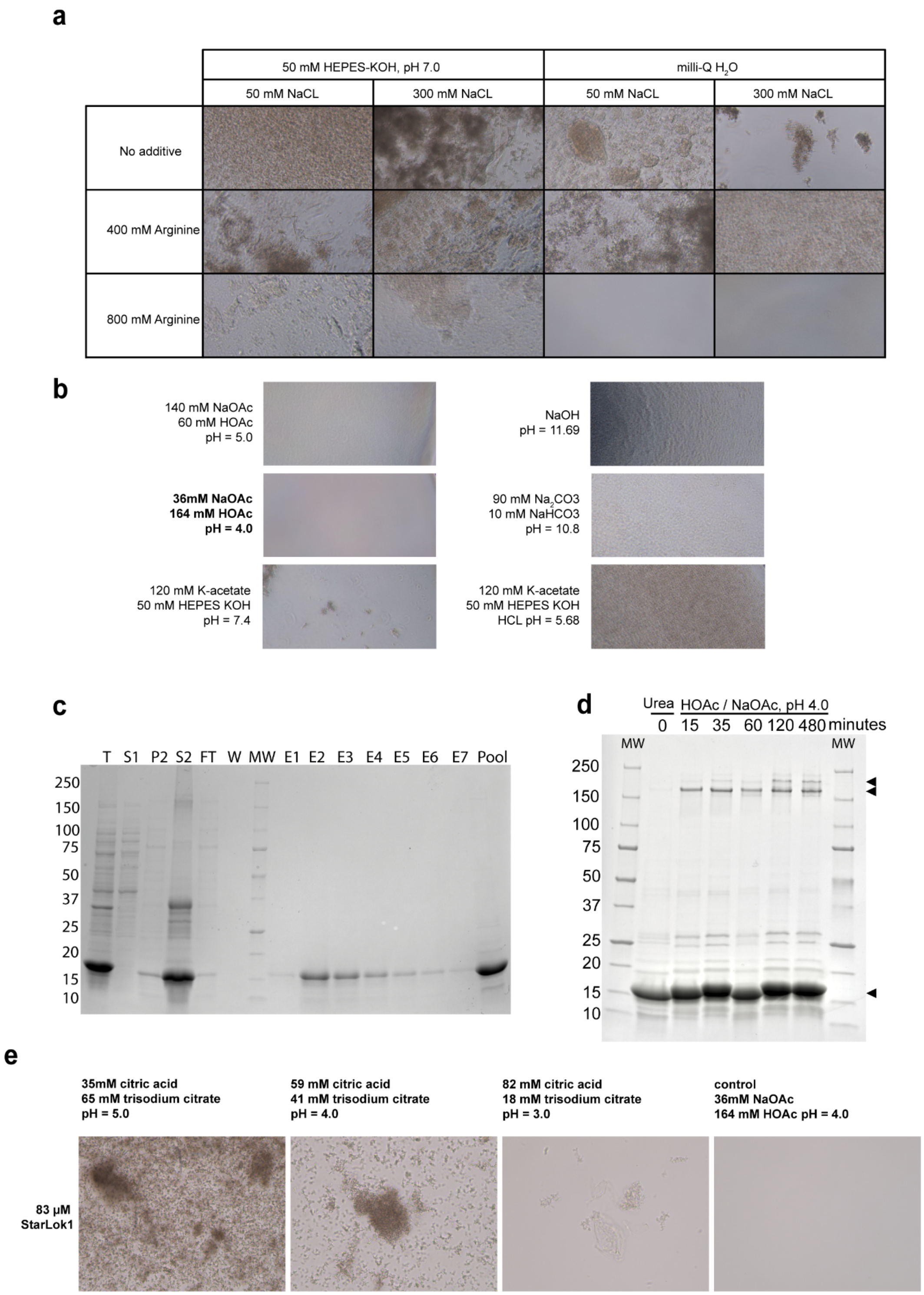


**Supplementary Fig. 2: Aggregation of concentrated StarLok1 is pH and solute dependent.** **a.** Light microscopy analysis of the StarLok1-containing sample after dialysis in buffers with different arginine and NaCl concentrations. Different levels of protein aggregation are observed in most conditions, but not in unbuffered solution of 800mM arginine. **b.** Light microscopy analysis of the StarLok1-containing sample after dialysis in buffers with different pH values and buffer compositions. At pH 4.0 maintained by a sodium acetate solution, StarLok1 does not show signs of aggregation unlike at higher pH. **c.** SDS-PAGE results of StarLok1 IMAC purification under denaturing condition with 6M urea. T: total, S1: supernatant before Urea addition, P2: pellet after urea denaturation, S2, supernatant after urea denaturation, FT: Flow through, W1 and W2: Washes, E1 to E7 corresponds to elution fractions. The pool fraction corresponds to the concentrated protein before refolding.  MW: Molecular weight marker. Numbering on the left indicates the apparent molecular weight in kDa. **d.** SDS-PAGE analysis of the kinetics of refolding of StarLok1. Arrow indicated the presence of StarLok1 as a monomer at the expected molecular weight of 18.6 kDa and the formation of multimers at an apparent molecular weight of ~ 180 kDa within the first 15 min of dialysis. **e.** Comparison of StarLok1 dialyzed in sodium citrate vs. sodium acetate. In acetate buffered solution, the protein is soluble at pH 4.0, but in citrate aggregates are still observed at pH 3.

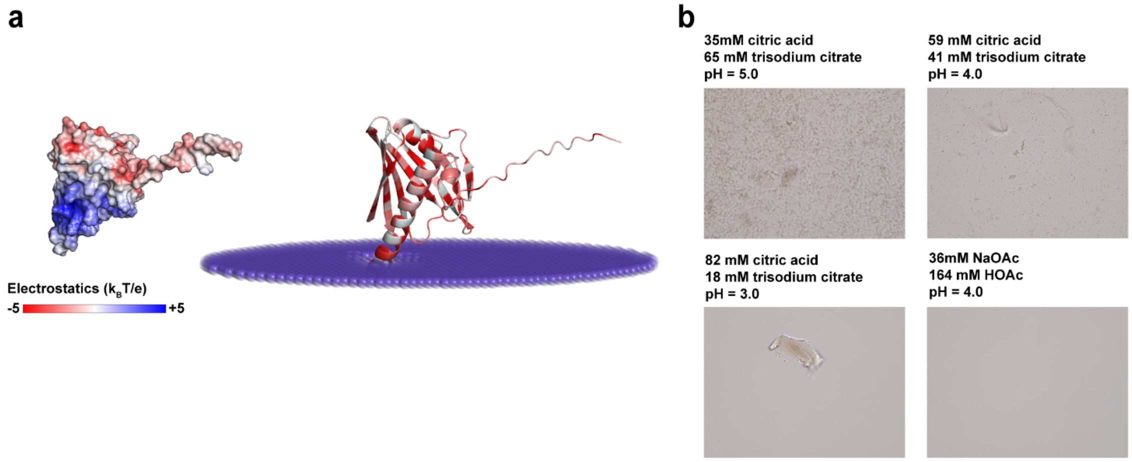


**Supplementary Fig. 3: Similarities between StarHod1 and StarLok1. a.** Structural prediction of StarHod1 as electrostatic surface potential ranging from -5 to +5 $k_B T/e$ and shown as a color gradient from red (negative) to white (neutral) to blue (positive) and corresponding membrane docking mode calculation on a model archaeal lipid membrane. The structure is colored according to the hydrophobicity of each residue from pale cyan (low hydrophobicity) to red (high hydrophobicity). **b.** Refolding of StarHod1 in different acidic buffers. Similarly to StarLok1, a combination of acetate and low pH is required for StarHod1 solubility.

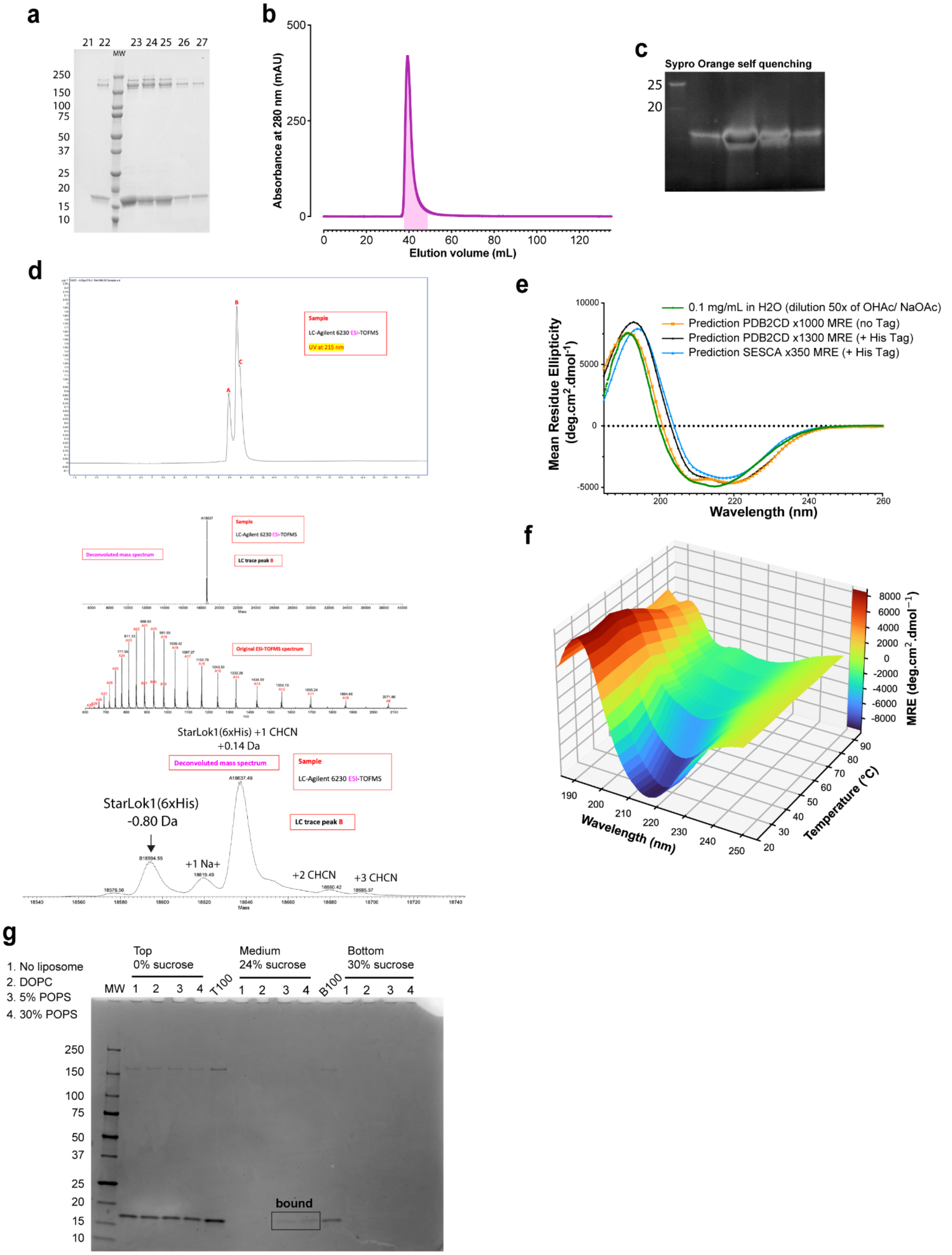


**Supplementary Fig. 4: Denaturation and refolding of StarLok1. a.** SDS-PAGE analysis from samples recovered from each fraction after size exclusion chromatography (fraction numbers are indicated at the top). MW: Molecular weight marker. Numbering on the left indicates the apparent molecular weight in kDa. **b.** SEC chromatogram, the plain pink area under the curve indicates the concentrated fraction (22 to 27) containing StarLok1. **c.** SDS-PAGE of the refolded StarLok1 in acetate buffer from fractions after size exclusion chromatography and stained Sypro orange. The dark spot in the second lane indicates oversaturation of the stain. **d.** LC-ESI-TOFMS analysis of recombinant StarLok1 (6xHis) purified from *E. coli i*n denatured condition and refolder in acetate buffer at pH 4.0. The chromatograms depicted the deconvoluted mass spectrum of the main detected peak. **e.** The CD spectrum of StarLok1 refolded in sodium acetate buffer is compared to theoretical CD spectra determined from Alphafold structures. **f.** Thermal denaturation of StarLok1 refolded in sodium acetate buffer and diluted in water as in panel E. The ellipticity of StarLok1 was monitored between 190 and 250 nm with increasing temperature from 20 to 90°C . The color gradient indicates MRE values and corresponding values are indicated on the right scale. As anticipated, a substantial decrease in β-sheet content at temperatures above 60°C is evidenced by the diminished characteristic negative ellipticity at 218 nm. In contrast, the α-helical structures exhibit greater thermal stability, maintaining their distinctive negative ellipticity peaks at 208 nm and 222 nm up to 90°C. This indicates that α-helical regions retain their secondary structure at elevated temperatures where β-sheet structures have predominantly denatured. **g.** Flotation assay of StarLok1 protein across sucrose density gradients following ultracentrifugation at 240,000 × *g* for 1 hr in the presence or absence of liposomes. SDS-PAGE analysis was used to determine the protein content in low-density (0% sucrose), intermediate-density (24% sucrose), and high-density (30% sucrose) fractions. StarLok1 was incubated with liposomes of varying compositions (samples 1–4) and homogenized in the high-density sucrose layer (30% w/v) before centrifugation. In the absence of anionic membranes, StarLok1 predominantly accumulates in the low-density fraction. However, when incubated with liposomes containing anionic membranes, the migration of StarLok1 to the low-density fraction is delayed, as evidenced by detectable amounts of the protein in the intermediate-density fraction (24% sucrose).

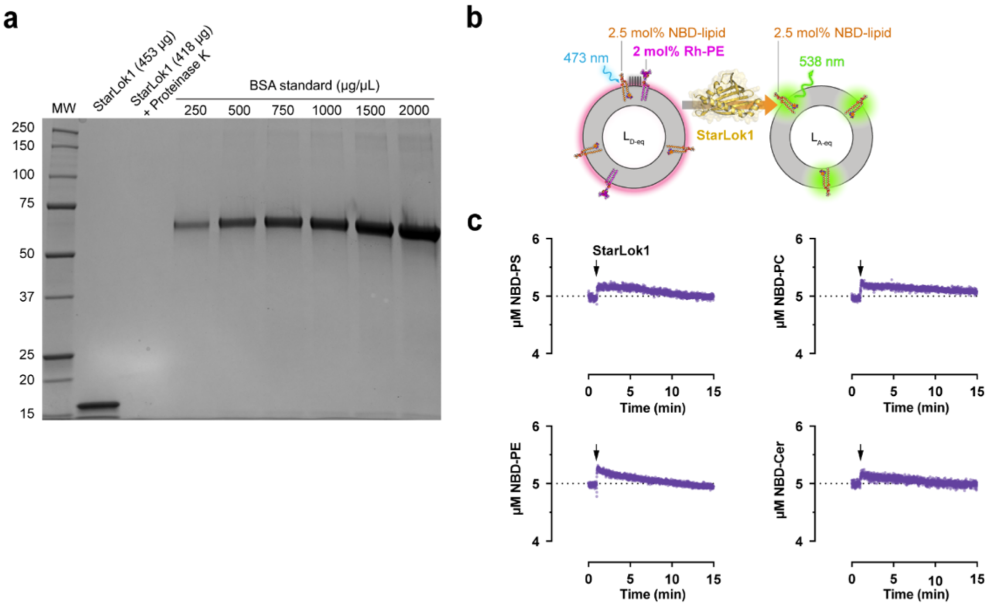


**Supplementary Fig. 5: Controls for StarLok1-mediated lipid transfer assay. a.** SDS–PAGE analysis of purified StarLok1 solubilized in N-lauroylsarcosine and used in the lipid transfer assay. Samples were analyzed before (lane 2) and after (lane 3) treatment with proteinase K. The amount of protein used in the LTP assay corresponds to approximately a 1:60 dilution of the sample loaded on the gel. BSA standards (lanes 4–9) were included to validate protein concentration determined by absorbance at 280 nm prior to the lipid transfer experiment. Molecular weight (MW) standard is indicated (lane 1). **b.** Schematic of the Rhodamine-PE transfer assay using donor ($L_{D\text{-eq}}$) and acceptor ($L_{A\text{-eq}}$) liposomes containing each 2.5 mol% NBD-lipids. The liposome composition shown represents the theoretical equilibrium state expected after complete LTP-mediated transfer of NBD-lipid (5 µM). In this configuration, transport of rhodamine-PE or liposome fusion, would result in decreased fluorescence due to FRET between NBD-lipid and rhodamine-PE in $L_{D\text{-eq}}$ liposomes. **c.** Kinetic trace of Rhodamine-PE transfer between $L_{D\text{-eq}}$ and $L_{A\text{-eq}}$ liposomes. Arrows indicate the injection of StarLok1. No appreciable Rhodamine-PE transport or liposome fusion was detected.

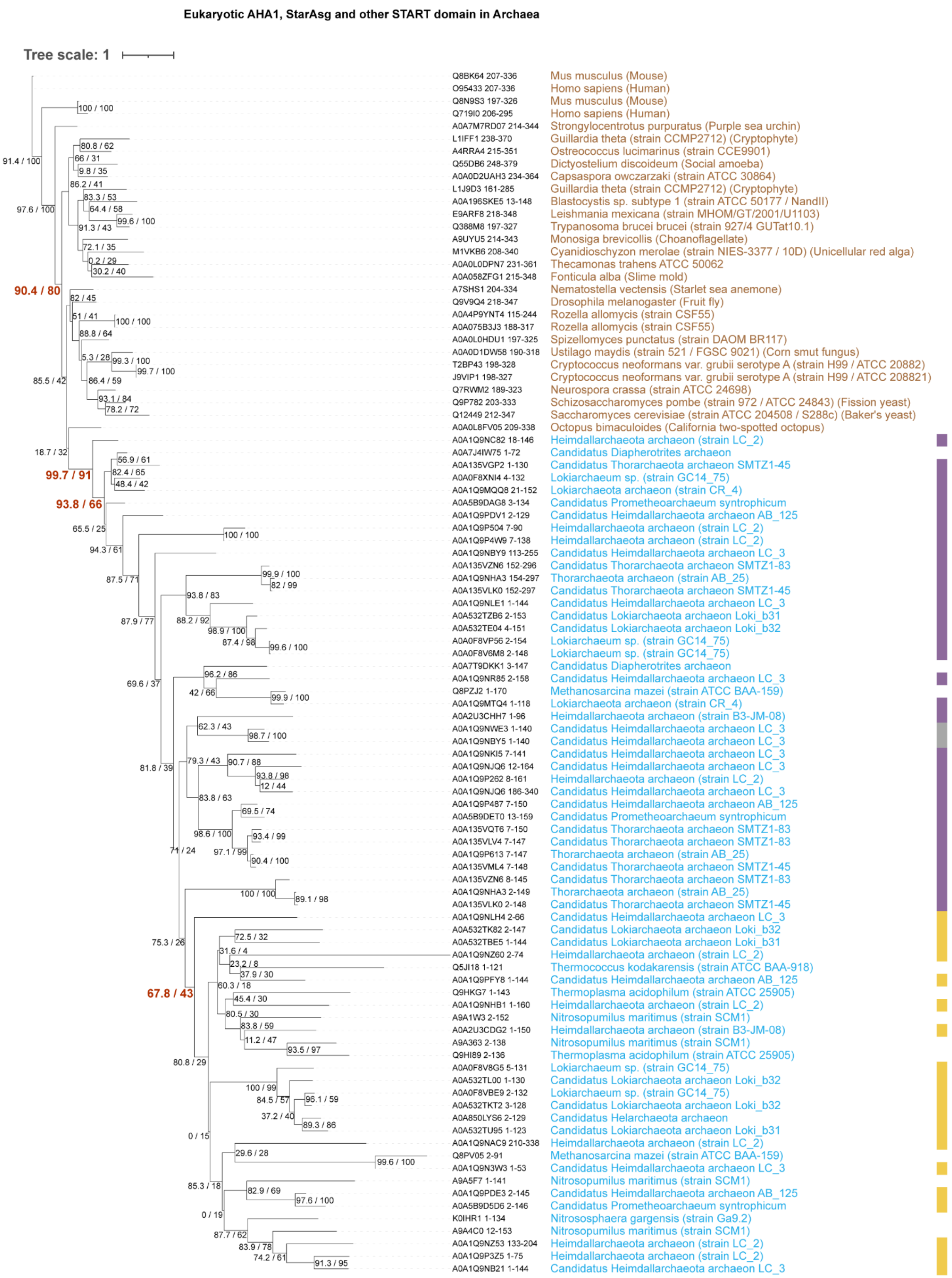

**Supplementary Fig. 6: Phylogenetic relationship between AHA1-like proteins and StarAsg proteins.** ML phylogenetic tree inferred from 96 START domain sequences. The dataset includes AHA1-like representatives from the following eukaryotic species: *Mus musculus, Homo sapiens,*

*Strongylocentrotus purpuratus, Guillardia theta, Ostreococcus lucimarinus, Dictyostelium discoideum, Capsaspora owczarzaki, Blastocystis sp., Leishmania mexicana, Trypanosoma brucei brucei, Monosiga brevicollis, Cyanidioschyzon merolae, Thecamonas trahens, Fonticula alba, Nematostella vectensis, Drosophila melanogaster, Rozella allomycis, Spizellomyces punctatus, Ustilago maydis, Cryptococcus neoformans, Neurospora crassa, Schizosaccharomyces pombe, Saccharomyces cerevisiae, and Octopus bimaculoides.* Archaeal sequences include all StarAsg1, StarAsg2, and StarAsg3, *Thermococcus kodakarensis, Thermoplasma acidophilum, Nitrosopumilus maritimus, Candidatus Diapherotrites archaeon, Methanosarcina mazei, and Nitrososphaera gargensis*. Sequences were aligned using MAFFT (G-INS-i strategy; --globalpair --maxiterate 1000; BLOSUM62 scoring matrix) with modified gap parameters (--op 1.53 --ep 0.5; --retree 2; --thread -1), generating alignment 297. The best-fit substitution model was selected using ModelFinder according to BIC (Q.PFAM+FO+G4), and ML inference was performed with IQ-TREE. Branch support values correspond to SH-aLRT (%) and ultrafast bootstrap (%) from 1,000 replicates each and are shown in parentheses. Yellow, purple, and gray stripes adjacent to branch-tip labels indicate Asgard sequences belonging to the StarAsg1, StarAsg2/3, or undefined clades, respectively (as in **Fig. 2a**). Branch lengths represent substitutions per site.

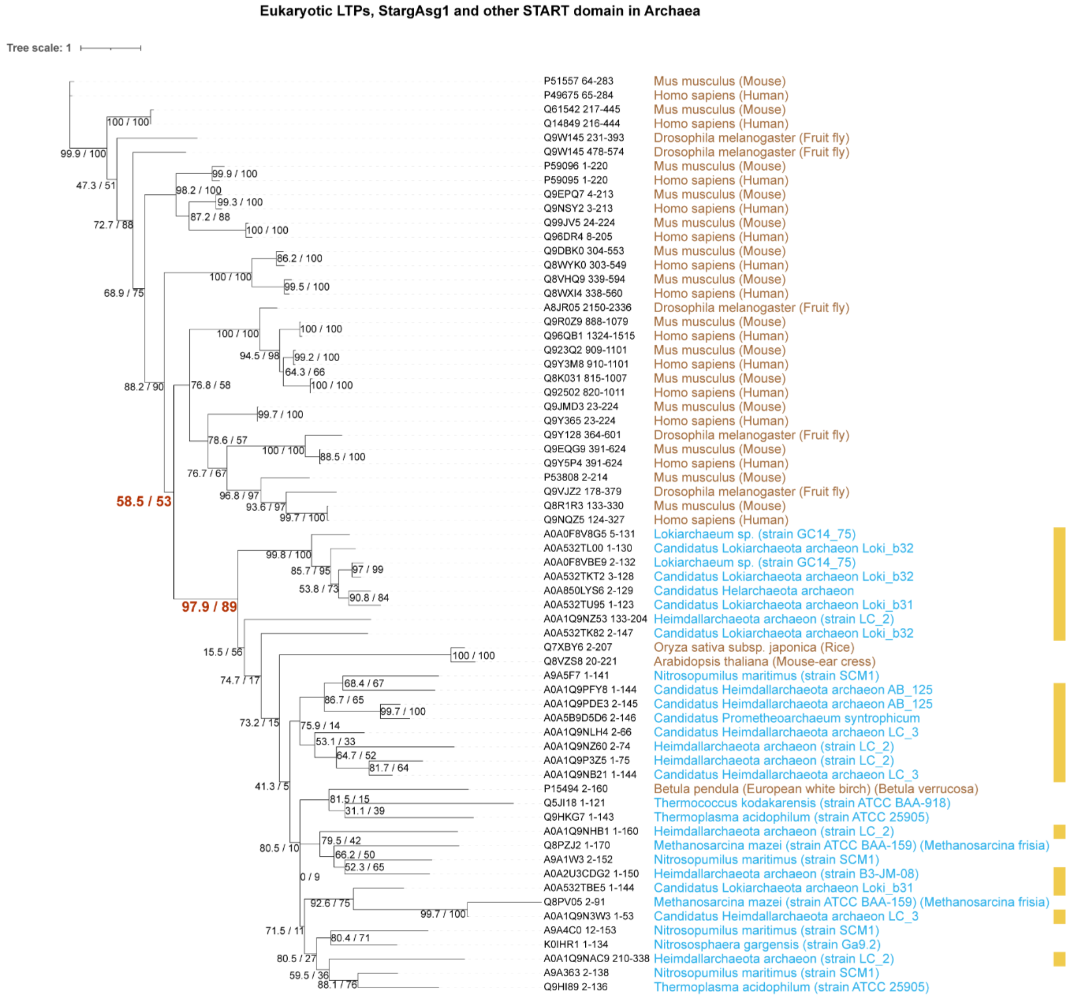


**Supplementary Fig. 7: Phylogenetic relationship between START LTPs and StarAsg1 proteins.** ML phylogenetic tree inferred from 65 START domain sequences. The dataset includes eukaryotic lipid transfer protein (LTP) representatives from Homo sapiens, Mus musculus, Drosophila melanogaster, Oryza sativa, Arabidopsis thaliana, and Betula pendula, together with StarAsg1 homologs from Asgard archaea and archaeal sequences from *Thermococcus kodakarensis, Thermoplasma acidophilum, Nitrosopumilus maritimus, Candidatus Diapherotrites archaeon, Methanosarcina mazei, and Nitrososphaera gargensis*. Sequences were aligned using MAFFT (L-INS-i strategy; --localpair --maxiterate 1000; BLOSUM62 scoring matrix) with modified gap parameters (--op 1.53 --ep 0.5; --retree 3; --thread -1), generating alignment 211. The best-fit substitution model was selected using ModelFinder according to BIC (Q.PFAM+FO+G4), and ML inference was performed with IQ-TREE. Branch support values correspond to SH-aLRT (%) and ultrafast bootstrap (%) from 1,000 replicates each and are shown in parentheses. Yellow stripes adjacent to branch-tip labels indicate StarAsg1 sequences. Branch lengths represent substitutions per site.

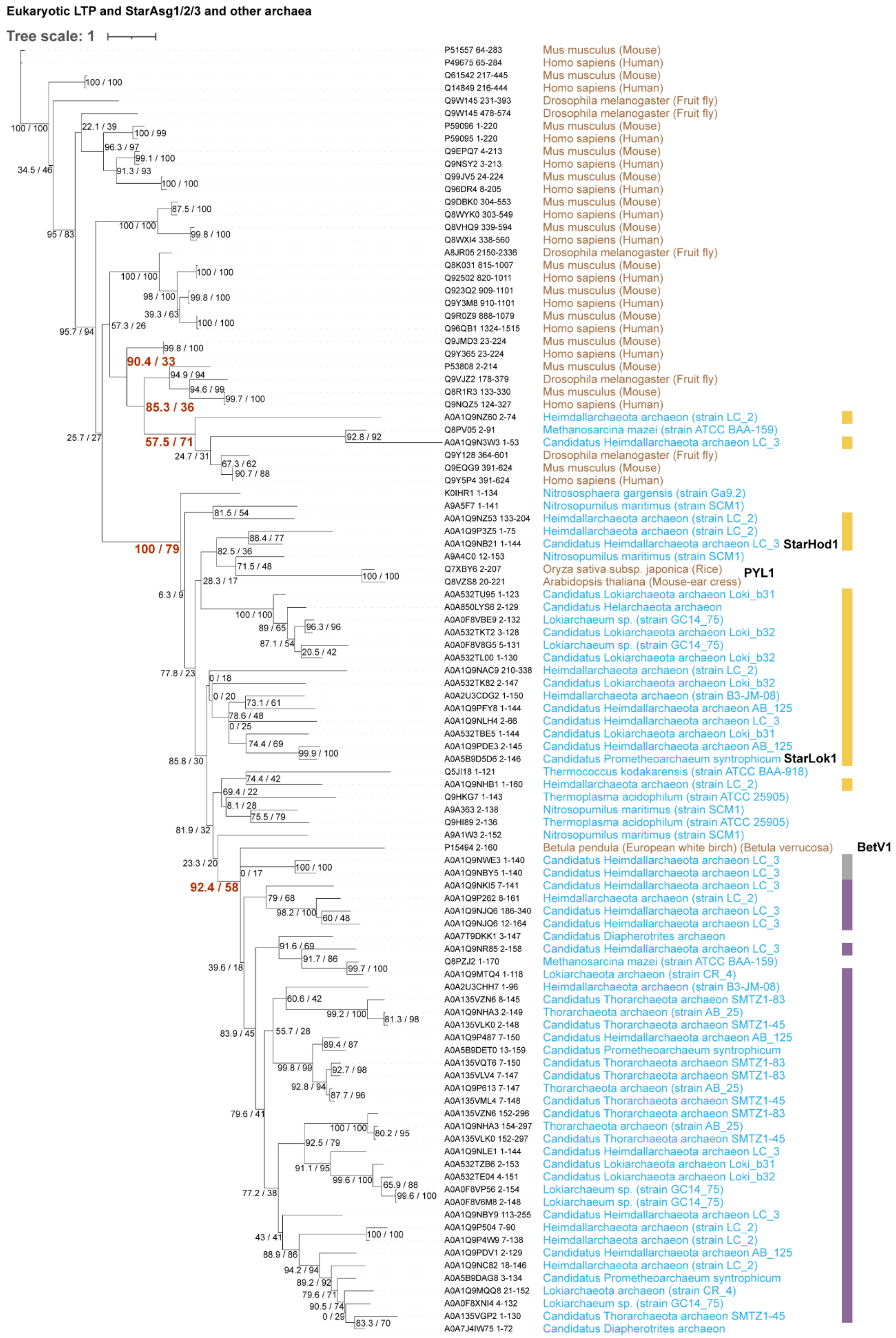


**Supplementary Fig. 8: Phylogenetic relationship between START LTPs and StarAsg proteins.** ML phylogenetic tree inferred from 102 START domain sequences. The dataset includes the same representative eukaryotic LTPs as described in **Supplementary Fig. 7** (*Homo*

*sapiens*, *Mus musculus*, *Drosophila melanogaster*, PYL1 from *Arabidopsis thaliana* and *Oryza sativa*, and Bet v 1 from *Betula pendula)*, excluding Eukaryotic AHA1-like protein. Sequences were aligned using MAFFT (L-INS-i strategy; --localpair --maxiterate 1000; BLOSUM80 scoring matrix) with modified gap parameters (--op 1.53 --ep 0.5; --retree 2). The best-fit substitution model was selected using ModelFinder according to BIC (Q.PFAM+FO+G4), and ML inference was performed with IQ-TREE. Branch support values correspond to SH-aLRT (%) and ultrafast bootstrap (%) from 1,000 replicates each and are shown in parentheses. Branch lengths represent substitutions per site. Branch supports of interest are highlighted in red. Yellow, purple, and gray stripes adjacent to branch-tip labels indicate Asgard sequences belonging to the StarAsg1, StarAsg2/3, or undefined clades, respectively (as in **Fig. 2a**). The tree is unrooted, although the outgroup taxon P49675 [65-284] (human STARD1) is displayed at the root for visualization purposes.

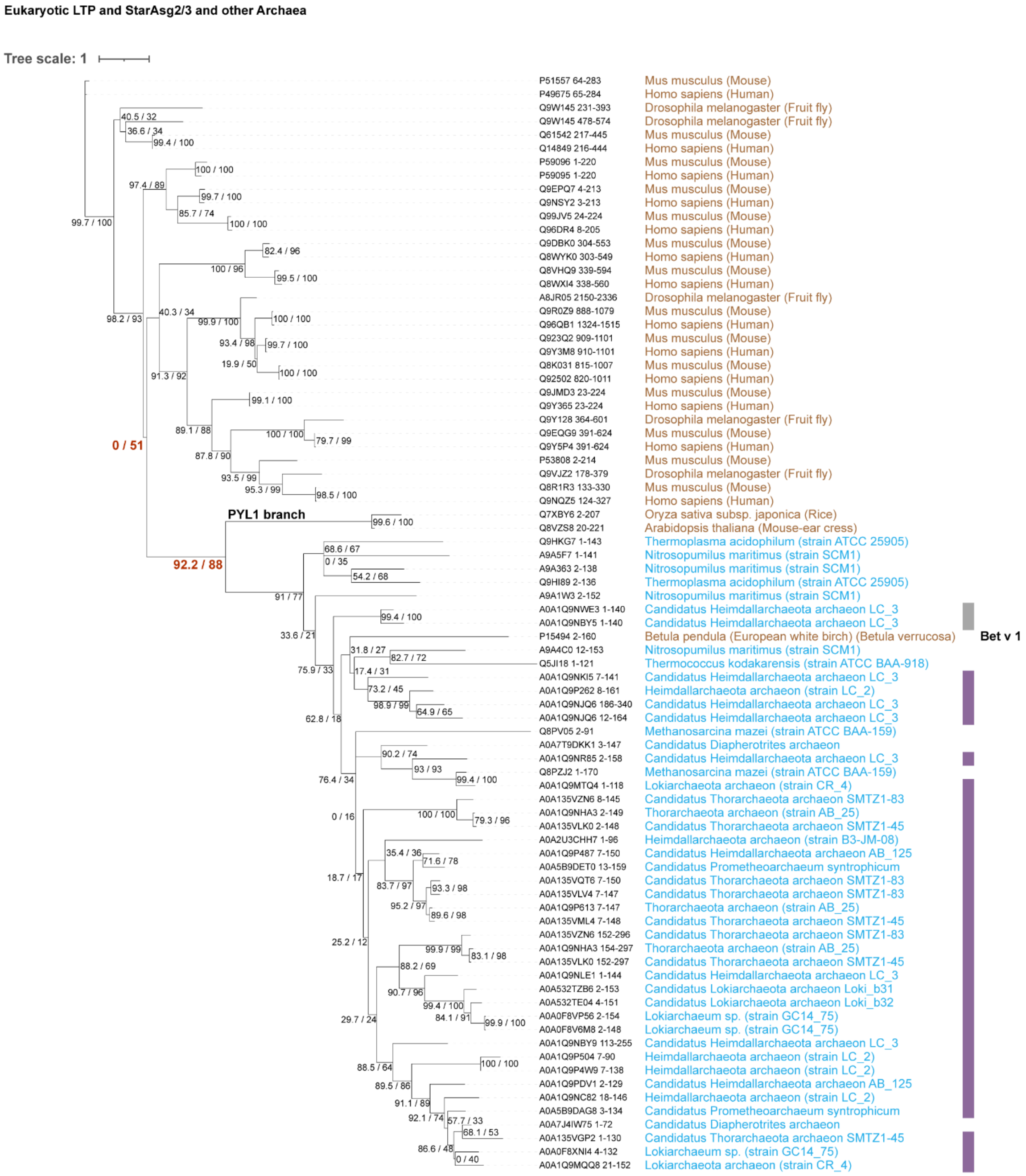


**Supplementary Fig. 9: Lack of phylogenetic relationship between START LTP and StarAsg2/3 proteins.** ML phylogenetic tree inferred from 81 START domain sequences. The dataset includes the same representative eukaryotic LTPs as described in **Supplementary Fig. 6** (*Homo sapiens*, *Mus musculus*, *Drosophila melanogaster*, PYL1 from *Arabidopsis thaliana* and *Oryza sativa*, and Bet v 1 from *Betula pendula)*, excluding Eukaryotic AHA1-like protein, and includes StarAsg2/3, and the archaeal species *Thermococcus kodakarensis*, *Thermoplasma acidophilum*, *Nitrosopumilus maritimus*, *Candidatus Diapherotrites archaeon*, *Methanosarcina*

*mazei*, *Nitrososphaera gargensis*. Sequences were aligned using MAFFT (L-INS-i strategy; --localpair --maxiterate 1000; BLOSUM80 scoring matrix) with modified gap parameters (--op 3.0 --ep 0.5 --retree 2 --lop -1.0 --lexp -0.5). The best-fit substitution model was selected using ModelFinder according to BIC (Q.PFAM+FO+G4), and ML inference was performed with IQ-TREE. Branch support values correspond to SH-aLRT (%) and ultrafast bootstrap (%) from 1,000 replicates each and are shown in parentheses. Branch supports of interest are highlighted in red. Purple and gray stripes adjacent to branch-tip labels indicate Asgard sequences belonging to StarAsg2/3 or undefined clades, respectively (as in **Fig. 2a**). Branch lengths represent substitutions per site.

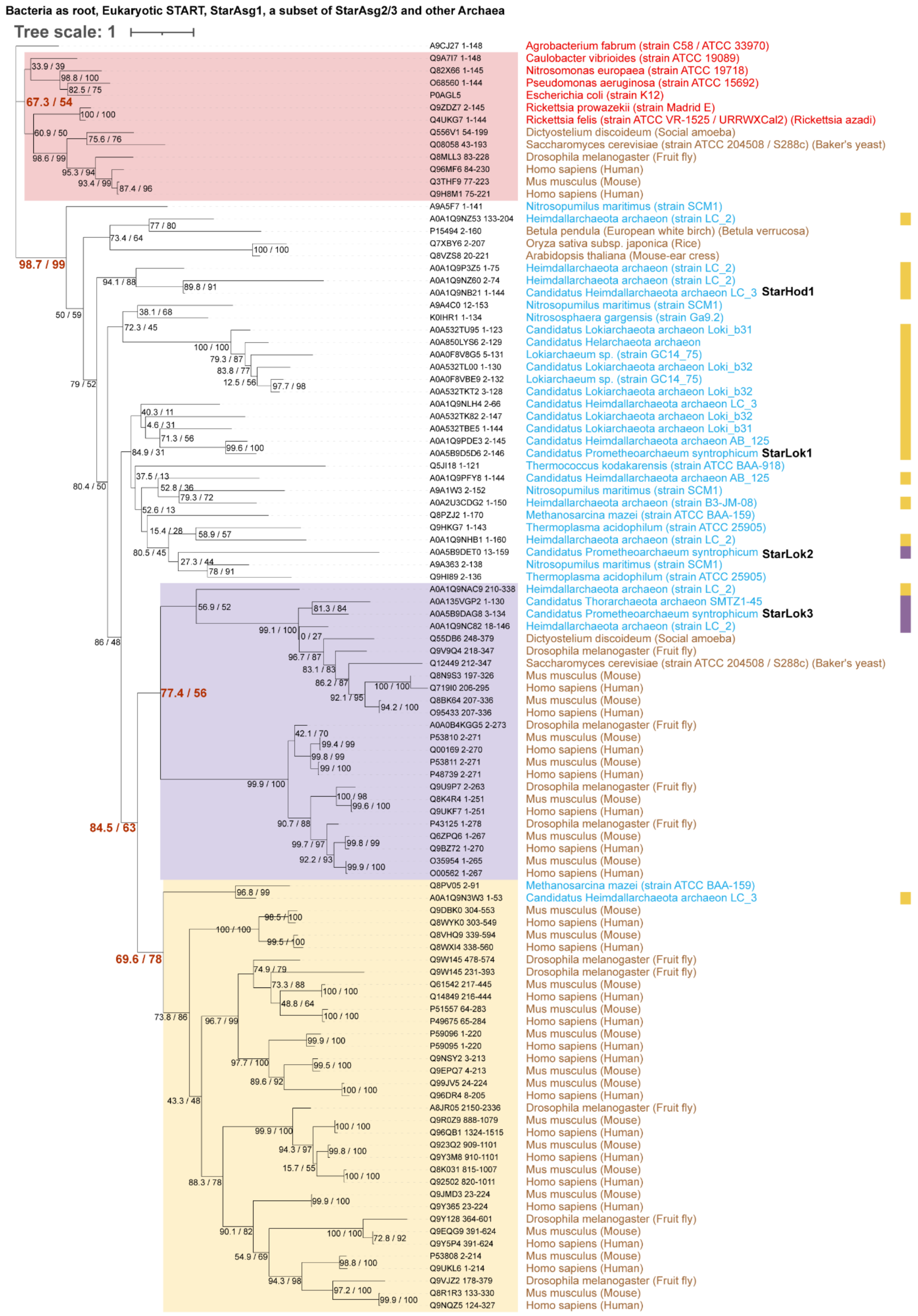


**Supplementary Fig. 10: Phylogenetic relationship of reduced START domain sample in the tree of life.** ML phylogenetic tree inferred from 103 START domain sequences. The dataset includes representative eukaryotes (*Dictyostelium discoideum, Homo sapiens, Mus musculus, Drosophila melanogaster, Saccharomyces cerevisiae, Betula pendula, Oryza sativa,* and *Arabidopsis thaliana*), encompassing AHA1-like representatives, canonical eukaryotic lipid transfer proteins (LTPs), and COQ-binding protein 10 homologs. The same archaeal species as in the previous datasets (**Supplementary Fig. 6**), however, only four StarAsg2/3 were retained, including sequences from *Promethearchaeum syntrophicum*. In addition, selected bacterial homologs were incorporated (*Agrobacterium fabrum, Caulobacter vibrioides, Nitrosomonas europaea, Pseudomonas aeruginosa, Escherichia coli, Rickettsia prowazekii*, and *Rickettsia felis*). Sequences were aligned using MAFFT (L-INS-i strategy; --localpair --maxiterate 1000; BLOSUM62) with modified gap parameters (--op 1.53 --ep 0.5 --retree 3 --lop -1.0 --lexp -0.5; --thread -1), generating alignment 212. The best-fit substitution model was selected using ModelFinder according to BIC (Q.PFAM+FO+G4), and ML inference was performed with IQ-TREE. Branch support values correspond to SH-aLRT (%) and ultrafast bootstrap (%) from 1,000 replicates each and are shown in parentheses. Yellow and Purple stripes adjacent to branch-tip labels indicate Asgard sequences belonging to StarAsg1 or StarAsg2/3, respectively (as in **Fig. 2a**). Branch lengths represent substitutions per site.

Batcteria (COQ10-like / RatA) vs Archaea (inc. StarAsg1/2/3)

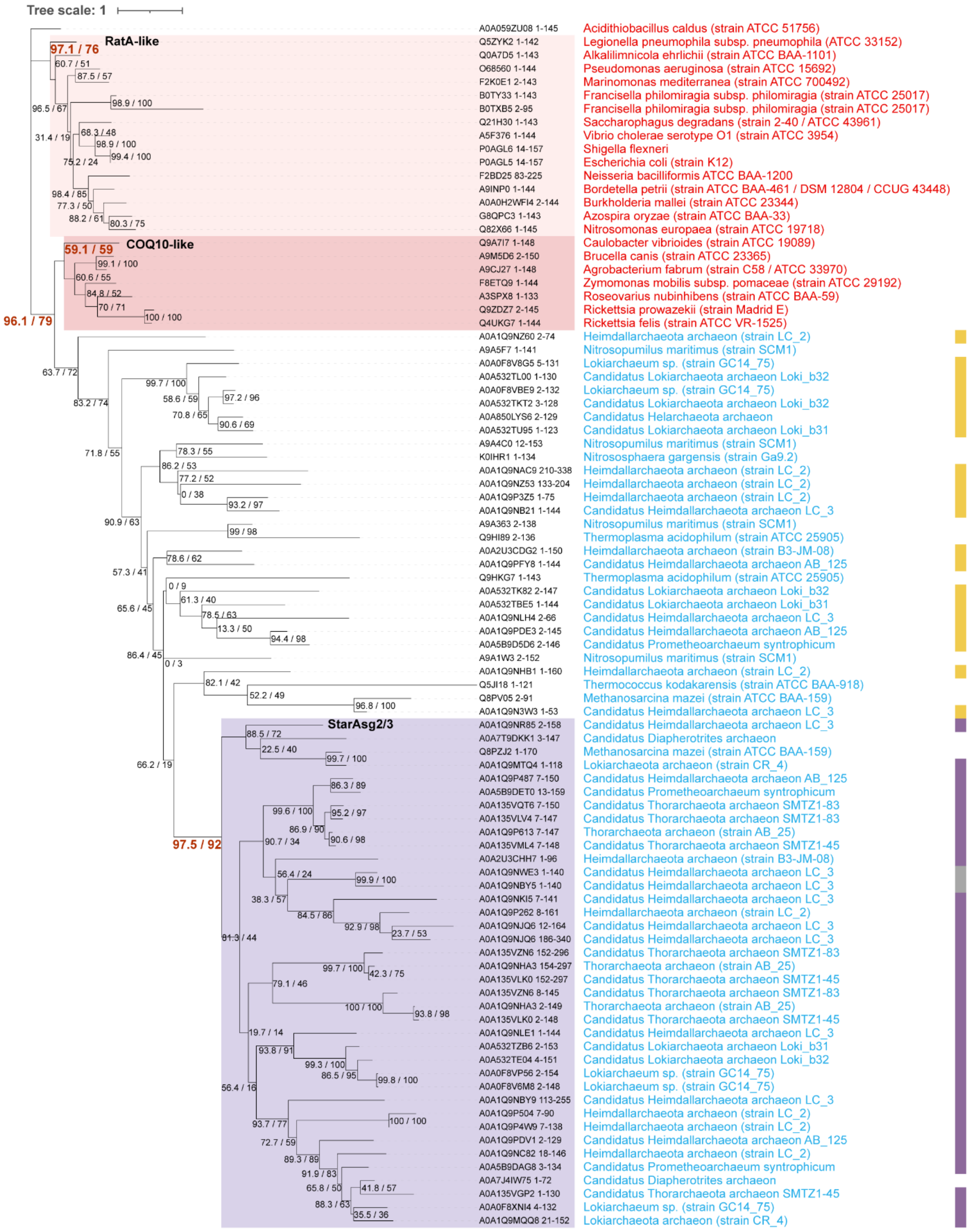


**Supplementary Fig. 11: Phylogenetic relationship bacterial RatA / COQ10 homologs with archaeal START domain**. ML phylogenetic tree inferred from 90 START domain sequences. The dataset includes the same archaeal species as in **Supplementary Fig. 6** and selected bacterial homologs (*Acidithiobacillus caldus, Legionella pneumophila subsp. pneumophila, Alkalilimnicola*

*ehrlichii, Pseudomonas aeruginosa, Marinomonas mediterranea, Francisella philomiragia subsp. philomiragia, Saccharophagus degradans, Vibrio cholerae, Shigella flexneri, Escherichia coli, Neisseria bacilliformis, Bordetella petrii, Burkholderia mallei, Azospira oryzae, Nitrosomonas europaea, Caulobacter vibrioides, Brucella canis, Agrobacterium fabrum, Zymomonas mobilis subsp. pomaceae, Roseovarius nubinhibens, Rickettsia prowazekii, and Rickettsia felis*). No eukaryotic sequences were included. Sequences were aligned using MAFFT (G-INS-i strategy; --globalpair --maxiterate 1000; BLOSUM62 scoring matrix) with modified gap parameters (--op 1.53 --ep 0.5; --retree 2; --thread -1), generating alignment 297. The best-fit substitution model was selected using ModelFinder according to BIC (Q.PFAM+FO+R4), and ML inference was performed with IQ-TREE. Branch support values correspond to SH-aLRT (%) and ultrafast bootstrap (%) from 1,000 replicates each and are shown in parentheses. Yellow, purple, and gray stripes adjacent to branch-tip labels indicate Asgard sequences belonging to the StarAsg1, StarAsg2/3, or undefined clades, respectively (as in **Fig. 2a**). Branch lengths represent substitutions per site.

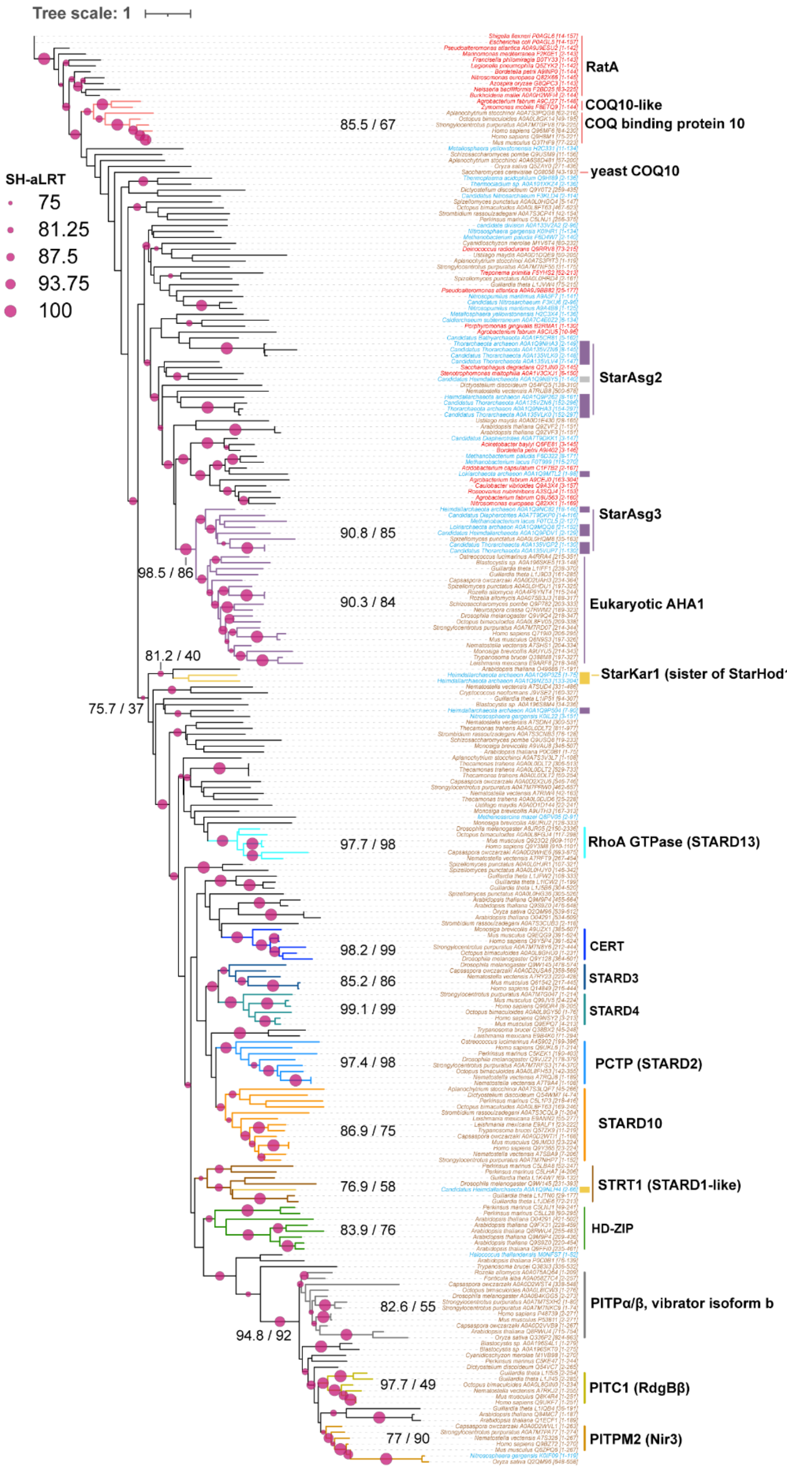


**Supplementary Fig. 12: Phylogenetic relationship START domain passing $\chi^2$ homogeneity composition test.** ML phylogenetic tree inferred from 247 START domain sequences that passed the χ² composition test. These sequences were selected from an initial dataset of 805 START domains extracted from 774 protein sequences spanning the tree of life. Sequences were aligned using MAFFT (E-INS-i strategy; --genafpair --maxiterate 1000; BLOSUM62 scoring matrix) with modified gap parameters (--op 1.53 --ep 0.123; --retree 3; --lop -1.0 --lexp -0.5; --LOP -8.00; --thread -1), generating alignment 24. The best-fit substitution model was selected using ModelFinder according to BIC (Q.PFAM+FO+R5), and ML inference was performed with IQ-TREE. Branch support values correspond to SH-aLRT (%) and ultrafast bootstrap (%) from 1,000 replicates each and are shown in parentheses. Yellow, purple, and gray stripes adjacent to branch-tip labels indicate Asgard sequences belonging to the StarAsg1, StarAsg2/3, or undefined clades, respectively (as in **Fig. 2a**). Branch lengths represent substitutions per site.

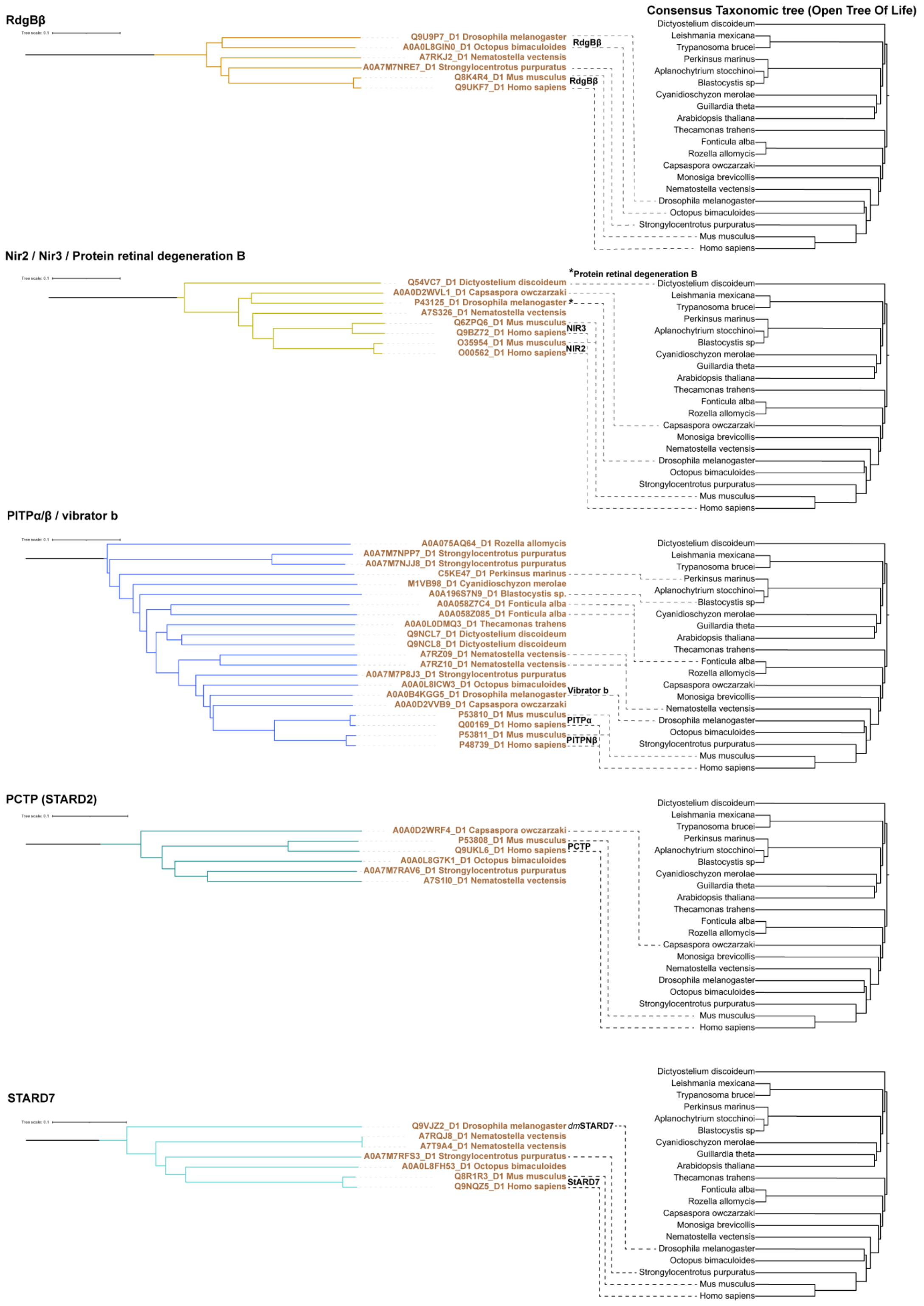


**Supplementary Fig. 13: Taxonomic congruence between eukaryotic START domain structural clades with Open Tree of Life taxonomy for RdgBβ, Nir2/Nir3, PITPα/β, PCTP (STARD2), and STARD7.** Comparison of START domain structural clades with species taxonomy derived from the Open Tree of Life (OTOL). For each START clade identified in the structural phylogeny, the corresponding subtree is shown on the left. The taxonomic tree inferred from OTOL for the same set of species is shown on the right. Protein domains are connected to their respective species when the distribution of sequences within a clade reflects the underlying species phylogeny, indicating taxonomic congruence. This panel includes the following START clades: RdgBβ (Nir2/Nir3; Protein retinal degeneration B), PITPα/β (phosphatidylinositol transfer proteins), PCTP (STARD2), and STARD7. Congruence is inferred qualitatively based on the correspondence between clade topology and accepted species relationships.

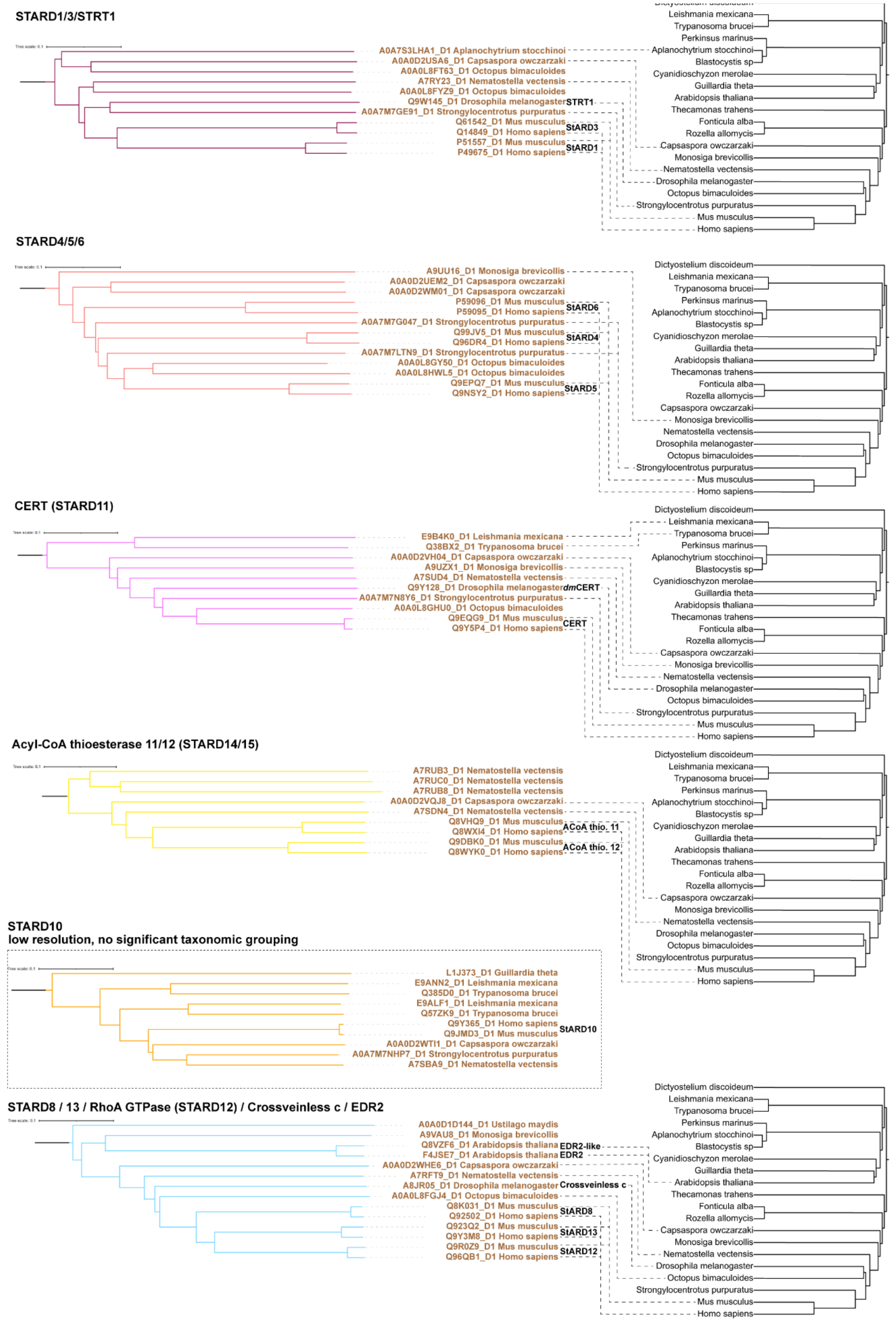


**Supplementary Fig. 14: Taxonomic congruence between eukaryotic START domain structural clades with Open Tree of Life taxonomy for STARD1/3, STARD4/5/6, CERT (STARD11), STARD14/15, STARD10, and STARD8/13/12-related proteins.** Comparison of

START domain structural clades with species taxonomy derived from the Open Tree of Life (OTOL). For each START clade identified in the structural phylogeny, the corresponding subtree is displayed on the left, and the OTOL taxonomic tree for the same species is shown on the right. Protein domains are linked with dashed lines to their respective species when the clade distribution mirrors species relationships, indicating taxonomic congruence. This panel includes the following START clades: STARD1/3, STARD4/5/6, CERT (STARD11), acyl-CoA thioesterase 11/12 (STARD14/15), STARD10, and STARD8/13/RhoA GTPase (STARD12)/Crossveinless c/EDR2. Most clades show a distribution broadly consistent with species phylogeny. In contrast, STARD10 exhibits limited taxonomic congruence and remains poorly resolved relative to accepted species relationships.

### Supplementary Text

Diversification of eukaryotic START-domain lipid transfer proteins

The broad taxon sampling of the structural phylogeny (**Fig. 5c**) resolves the eukaryotic diversification of START LTPs into previously described subclasses[1–3], summarized in the cladogram of Figure 5d. Because FoldTree does not provide node-level statistical support, we assessed the reliability of these groupings through their taxonomic congruence with the Open Tree of Life (**Supplementary Figs. 13, 14**). Where the reduced ML dataset contains the relevant taxa, the same groupings are recovered in the ML phylogeny (**Fig. 5a**), providing independent corroboration.

Recovered relationships between eukaryotic START LTPs include orthology between STARD4/5/6 and STARD1/3, both of which radiate independently in animals and their closest unicellular relatives, such as *Capsaspora owczarzaki*, and which are both recovered in the ML phylogeny (SH-aLRT/UFBoot = 90.6/87 and 97.1/98, respectively; **Fig. 5a**). Similarly, the two START-domain-containing acyl-CoA thioesterases (STARD14 and STARD15) show orthology, also recovered in the ML tree (93.2/84), and arose prior to the divergence of animals and *C. owczarzaki*. More distantly, CERT (STARD11) groups as an independent class. We also observed orthology among STARD8, STARD12, and STARD13, which contain a START domain fused to a RhoGAP domain, are common to choanoflagellates, and group in the structural phylogeny with crossveinless c and the plant EDR2-like proteins; this clade is recovered in the ML tree (97.8/100) though plant sequences are absent from that dataset. Our trees also suggest that STARD2 (PCTP) and STARD7 form a paralogous sister group relative to other STARD members, both recovered in the ML phylogeny (84.1/82 and 100/100). STARD10 was the only subclass showing poor taxonomic congruence with the Open Tree of Life and is therefore treated as unresolved. The PITP class, which transports phosphatidylinositol and phosphatidic acid[4,5], consistently diverged from STARD proteins at the earliest stage of START LTP radiation in eukaryotes, supporting the acquisition of phosphatidylinositol as a key lipid in eukaryotes.
